# Whole genome sequencing of a sporadic primary immunodeficiency cohort

**DOI:** 10.1101/499988

**Authors:** James E. D. Thaventhiran, Hana Lango Allen, Oliver S. Burren, William Rae, Daniel Greene, Emily Staples, Zinan Zhang, James H. R. Farmery, Ilenia Simeoni, Elizabeth Rivers, Jesmeen Maimaris, Christopher J Penkett, Jonathan Stephens, Sri V.V. Deevi, Alba Sanchis-Juan, Nicholas S Gleadall, Moira J. Thomas, Ravishankar B. Sargur, Pavels Gordins, Helen E. Baxendale, Matthew Brown, Paul Tuijnenburg, Austen Worth, Steven Hanson, Rachel Linger, Matthew S. Buckland, Paula J. Rayner-Matthews, Kimberly C. Gilmour, Crina Samarghitean, Suranjith L. Seneviratne, David M. Sansom, Andy G. Lynch, Karyn Megy, Eva Ellinghaus, David Ellinghaus, Silje F. Jorgensen, Tom H Karlsen, Kathleen E. Stirrups, Antony J. Cutler, Dinakantha S. Kumararatne, Anita Chandra, J. David M. Edgar, Archana Herwadkar, Nichola Cooper, Sofia Grigoriadou, Aarnoud Huissoon, Sarah Goddard, Stephen Jolles, Catharina Schuetz, Felix Boschann, NBR-RD PID Consortium, NIHR BioResource, Paul A. Lyons, Matthew E. Hurles, Sinisa Savic, Siobhan O. Burns, Taco W. Kuijpers, Ernest Turro, Willem H. Ouwehand, Adrian J. Thrasher, Kenneth G. C. Smith

**Affiliations:** Cambridge Institute of Therapeutic Immunology and Infectious Disease, Jeffrey Cheah Biomedical Centre, Cambridge Biomedical Campus, Cambridge, UK; Department of Medicine, University of Cambridge School of Clinical Medicine, Cambridge Biomedical Campus, Cambridge, UK; Medical Research Council Toxicology Unit, School of Biological Sciences, University of Cambridge, Cambridge, UK; Department of Haematology, University of Cambridge School of Clinical Medicine, Cambridge Biomedical Campus, Cambridge, UK; NHS Blood and Transplant, Cambridge Biomedical Campus, Cambridge, UK; NIHR BioResource, Cambridge University Hospitals, Cambridge Biomedical Campus, Cambridge, UK; Medical Research Council Epidemiology Unit, University of Cambridge School of Clinical Medicine, Cambridge Biomedical Campus, Cambridge, UK; Medical Research Council Biostatistics Unit, Cambridge Biomedical Campus, Cambridge, UK; Molecular Development of the Immune System Section, Laboratory of Immune System Biology and Clinical Genomics Program, National Institute of Allergy and Infectious Diseases, National Institutes of Health, Bethesda, MD, USA; Cancer Research UK Cambridge Institute, University of Cambridge, Li Ka Shing Centre, Robinson Way, Cambridge, UK; UCL Great Ormond Street Institute of Child Health, London, UK; Great Ormond Street Hospital for Children NHS Foundation Trust, London, UK; Department of Immunology, Queen Elizabeth University Hospital, Glasgow, UK; Sheffield Teaching Hospitals NHS Foundation Trust, Sheffield, UK; Department of Infection Immunity and Cardiovascular Disease, University of Sheffield, Sheffield, UK; East Yorkshire Regional Adult Immunology and Allergy Unit, Hull Royal Infirmary, Hull and East Yorkshire Hospitals NHS Trust, Hull, UK; Royal Papworth Hospital NHS Foundation Trust, Cambridge, UK; Department of Pediatric Immunology, Rheumatology and Infectious Diseases, Emma Children’s Hospital & The Department of Experimental Immunology, Amsterdam University Medical Center (AMC), University of Amsterdam, Amsterdam, The Netherlands; Institute of Immunity and Transplantation, University College London, London, UK; Department of Immunology, Royal Free London NHS Foundation Trust, London, UK; School of Mathematics and Statistics/School of Medicine, University of St Andrews, St Andrews, UK; K.G. Jebsen Inflammation Research Centre, Institute of Clinical Medicine, University of Oslo, Oslo University Hospital, Rikshospitalet, Oslo, Norway; Department of Transplantation, Institute of Clinical Medicine, University of Oslo, Oslo University Hospital, Rikshospitalet, Oslo, Norway; Institute of Clinical Molecular Biology, Christian Albrechts University of Kiel, Kiel, Germany; Section of Clinical Immunology and Infectious Diseases, Department of Rheumatology, Dermatology and Infectious Diseases, Oslo University Hospital, Rikshospitalet, Norway; Research Institute of Internal Medicine, Division of Surgery, Inflammatory Diseases and Transplantation, Oslo University Hospital, Rikshospitalet, Norway; JDRF/Wellcome Diabetes and Inflammation Laboratory, Wellcome Centre for Human Genetics, Nuffield Department of Medicine, NIHR Oxford Biomedical Research Centre, University of Oxford, Oxford, UK; Department of Clinical Biochemistry and Immunology, Cambridge University Hospitals, Cambridge Biomedical Campus, Cambridge, UK; St James’s Hospital & Trinity College Dublin, Ireland; Salford Royal NHS Foundation Trust, Salford, UK; Department of Medicine, Imperial College London, London, UK; Barts Health NHS Foundation Trust, London, UK; West Midlands Immunodeficiency Centre, University Hospitals Birmingham, Birmingham, UK; University Hospitals of North Midlands NHS Trust, Stoke-on-Trent, UK; Immunodeficiency Centre for Wales, University Hospital of Wales, Cardiff, UK; Department of Pediatric Immunology, University Hospital Carl Gustav Carus, Dresden, Germany; Institute of Medical Genetics and Human Genetics, Charite-Universitatsmedizin Berlin, Berlin, Germany; Department of Human Genetics, Wellcome Sanger Institute, Wellcome Genome Campus, Hinxton, Cambridge, UK; The Department of Clinical Immunology and Allergy, St James’s University Hospital, Leeds, UK; The NIHR Leeds Biomedical Research Centre and Leeds Institute of Rheumatic and Musculoskeletal Medicine, Leeds, UK

## Abstract

Primary immunodeficiency (PID) is characterised by recurrent and often life-threatening infections, autoimmunity and cancer, and it presents major diagnostic and therapeutic challenges. Although the most severe forms present in early childhood, the majority of patients present in adulthood, typically with no apparent family history and a variable clinical phenotype of widespread immune dysregulation: about 25% of patients have autoimmune disease, allergy is prevalent, and up to 10% develop lymphoid malignancies^1–3^. Consequently, in sporadic PID genetic diagnosis is difficult and the role of genetics is not well defined. We addressed these challenges by performing whole genome sequencing (WGS) of a large PID cohort of 1,318 participants. Analysis of coding regions of 886 index cases found disease-causing mutations in known monogenic PID genes in 10.3%, while a Bayesian approach (BeviMed^4^) identified multiple potential new candidate genes, including *IVNS1ABP*. Exploration of the non-coding genome revealed deletions in regulatory regions which contribute to disease causation. Finally, a genome-wide association study (GWAS) identified PID-associated loci and uncovered evidence for co-localisation of, and interplay between, novel high penetrance monogenic variants and common variants (at the *PTPN2* and *SOCS1* loci). This begins to explain the contribution of common variants to variable penetrance and phenotypic complexity in PID. Thus, a cohort-based WGS approach to PID diagnosis can increase diagnostic yield while deepening our understanding of the key pathways influencing human immune responsiveness.

---

The phenotypic heterogeneity of PID leads to diagnostic difficulty, and almost certainly to an underestimation of its true incidence. Our cohort reflects this heterogeneity, though it is dominated by adult onset, sporadic antibody deficiency-associated PID (AD-PID: comprising Common Variable Immunodeficiency (CVID), Combined Immunodeficiency (CID) and isolated antibody deficiency). Identifying a specific genetic cause of PID can facilitate definitive treatment including haematopoietic stem cell transplantation, genetic counselling, and the possibility of gene-specific therapy^2^ while contributing to our understanding of the human immune system^5^. Unfortunately, only 29% of patients with PID have a genetic cause of their disease identified^6^, with the lowest rate in patients who present as adults and have no apparent family history. While variants in over 300 genes have been described as monogenic causes of PID^3^, it is often difficult to match the clinical phenotype to a known genetic cause, because phenotypes are heterogeneous and disease penetrance is often low^2,7^. Furthermore, a common variant analysis of CVID identified new disease-associated loci, and raised the possibility that common variants may impact upon clinical presentation^8^. We therefore investigated whether applying WGS across a “real world” PID cohort might illuminate the complex genetics of the range of conditions collectively termed PID: the approach is summarised in **Extended Data Fig. 1**.

## Patient cohort

We sequenced 1,318 individuals recruited as part of the PID domain of the United Kingdom NIHR BioResource - Rare Diseases program (NBR-RD; **Extended Data Fig.2; Supplementary Methods**). The cohort comprised of both sporadic and familial PID patients (N=974) and family members. Of the patients, 886 were index cases who fell into one of the diagnostic categories of the European Society for Immunodeficiencies (ESID) registry diagnostic criteria (**Fig. 1a; Extended Data Table 1**). This cohort represents a third of CVID and half of CID patients registered in the UK^9^. Clinical phenotypes were dominated by adult-onset sporadic AD-PID: all had recurrent infections, 28% had autoimmunity, and 8% had malignancy (**Fig. 1a-b, Extended Data Table 2**), mirroring the UK national PID registry^6^.

**Figure 1.**
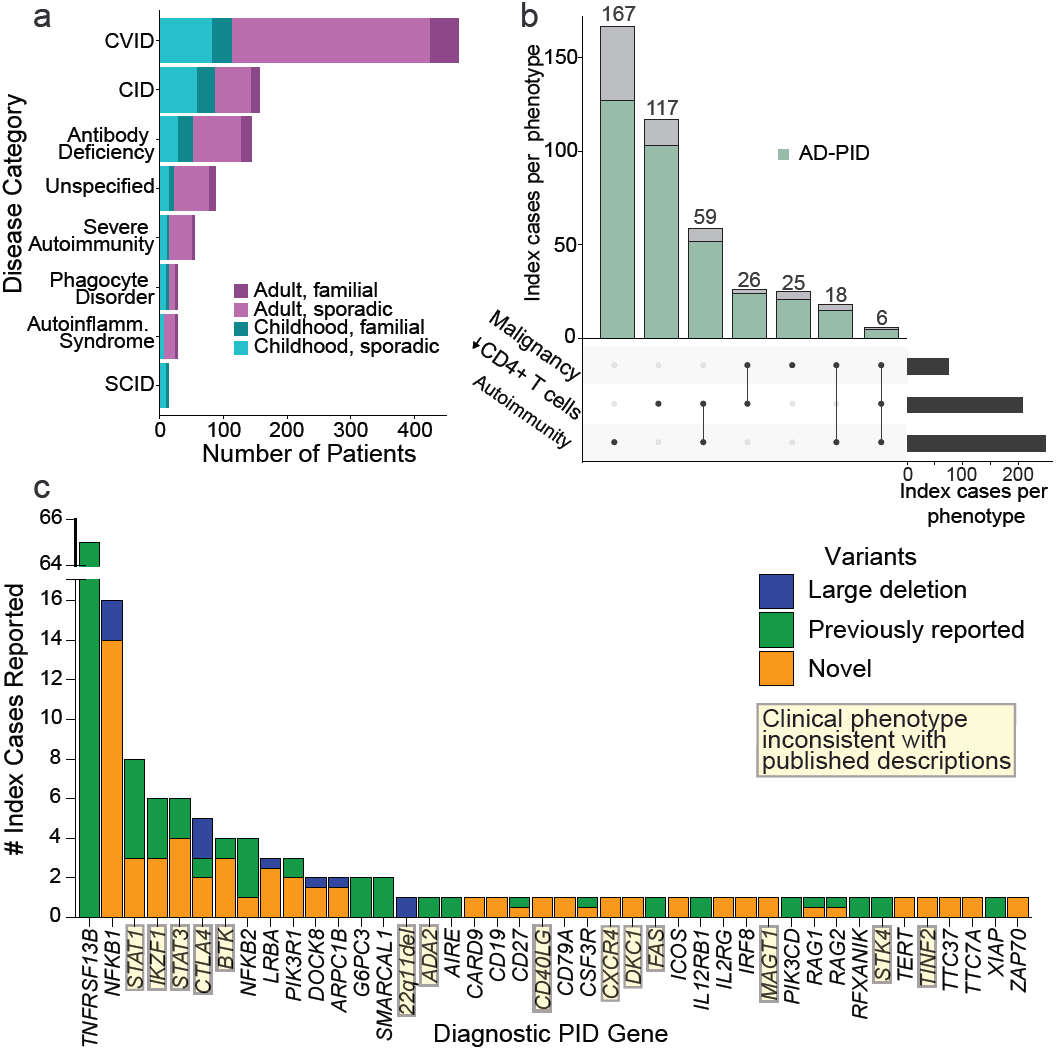
Description of the immunodeficiency cohort and disease associations in coding regions. **(a)** Number of index cases recruited under different phenotypic categories (red – adult cases, blue – paediatric cases, lighter shade – sporadic (no family history of PID), darker shade - family history of PID). CVID – Common variable immunodeficiency, CID – combined immunodeficiency, and SCID – severe combined immunodeficiency. **(b)** Number of index cases with malignancy, autoimmunity and CD4+ lymphopenia. (black bar – total number of cases, blue bar - number of cases with AD-PID phenotype). **(c)** Number of patients with reported genetic findings subdivided by gene. Previously reported variants are those identified as immune disease-causing in the HGMD-Pro database.

## Identification of Pathogenic Variants in Known Genes

We analysed coding regions of genes with previously reported disease-causing variants in PID^10^ (**Methods**). Based on filtering criteria for diagnostic reporting according to the American College of Medical Genetics (ACMG) guidelines^11^ and described in the Methods, we identified and reported to the referring clinicians 104 known or likely pathogenic variants in 91 index cases (10.3%) across 41 genes implicated in monogenic disease (**Fig. 1c; Supplementary Table 1**). 60 patients (6.8%) had a previously reported pathogenic variant in the disease modifier *TNFRSF13B* (*TACI*), increasing the proportion of cases with a reportable variant to 17.0% (151 patients). Interestingly, 5 patients with a monogenic diagnosis (in *BTK, LRBA, MAGT1, RAG2, SMARCAL1*) also had a pathogenic *TNFRSF13B* variant. Of the 103 monogenic variants we report here, 69 (67.0%) had not been previously described (**Supplementary Table 1**) and 8 were structural variants, including single exon and non-coding promoter deletions unlikely to have been detected by whole exome sequencing^12^.

In 22 patients with variants in 14 genes (34% of 41 identified genes) reported as pathogenic, the clinical presentation deviated from the phenotypes typically associated with those genes. One example was chronic mucocutaneous candidiasis (CMC), which is the trigger for clinical genetic testing for *STAT1* GOF variants, as CMC was reported in 98% of such patients^13,14^. Now this series, along with single case reports^15,16^, indicate *STAT1* GOF may present with phenotypes as diverse as CVID or primary antibody deficiency. Since many PID-associated genes were initially discovered in a small number of familial cases, it is not surprising that the phenotypes described in the literature do not reflect the true clinical diversity. Thus, a cohort-based WGS approach to PID provides a diagnostic yield even in a predominantly sporadic cohort, allows diagnoses which are not constrained by pre-existing assumptions about genotype-phenotype relationships, and suggests caution in the use of clinical phenotype in targeted gene screening and interpreting PID genetic data.

## An approach to prioritising candidate PID-associated genes in a WGS cohort

We next determined whether the cohort-based WGS approach could identify new genetic associations with PID. We included all 886 index cases in a single cohort in order to optimise statistical power, and because genotype-phenotype correlation in PID is incompletely understood. We applied a Bayesian inference procedure, named BeviMed^4^, and used it to determine posterior probabilities of association (PPA) between each gene and case/control status of the 886 index cases and 9,283 unrelated controls (**Methods**). We obtained a BeviMed PPA for 31,350 genes in the human genome; the 25 highest ranked genes are shown in **Fig. 2a** (see also **Supplementary Table 2** and **Supplementary Note 2**). Overall, genes with BeviMed PPA>0.1 were strongly enriched for known PID genes (odds ratio = 15.1, P = 3.1×10^−8^ Fisher’s Exact test), demonstrating that a statistical genetic association approach can identify genes causal for PID.

**Figure 2.**
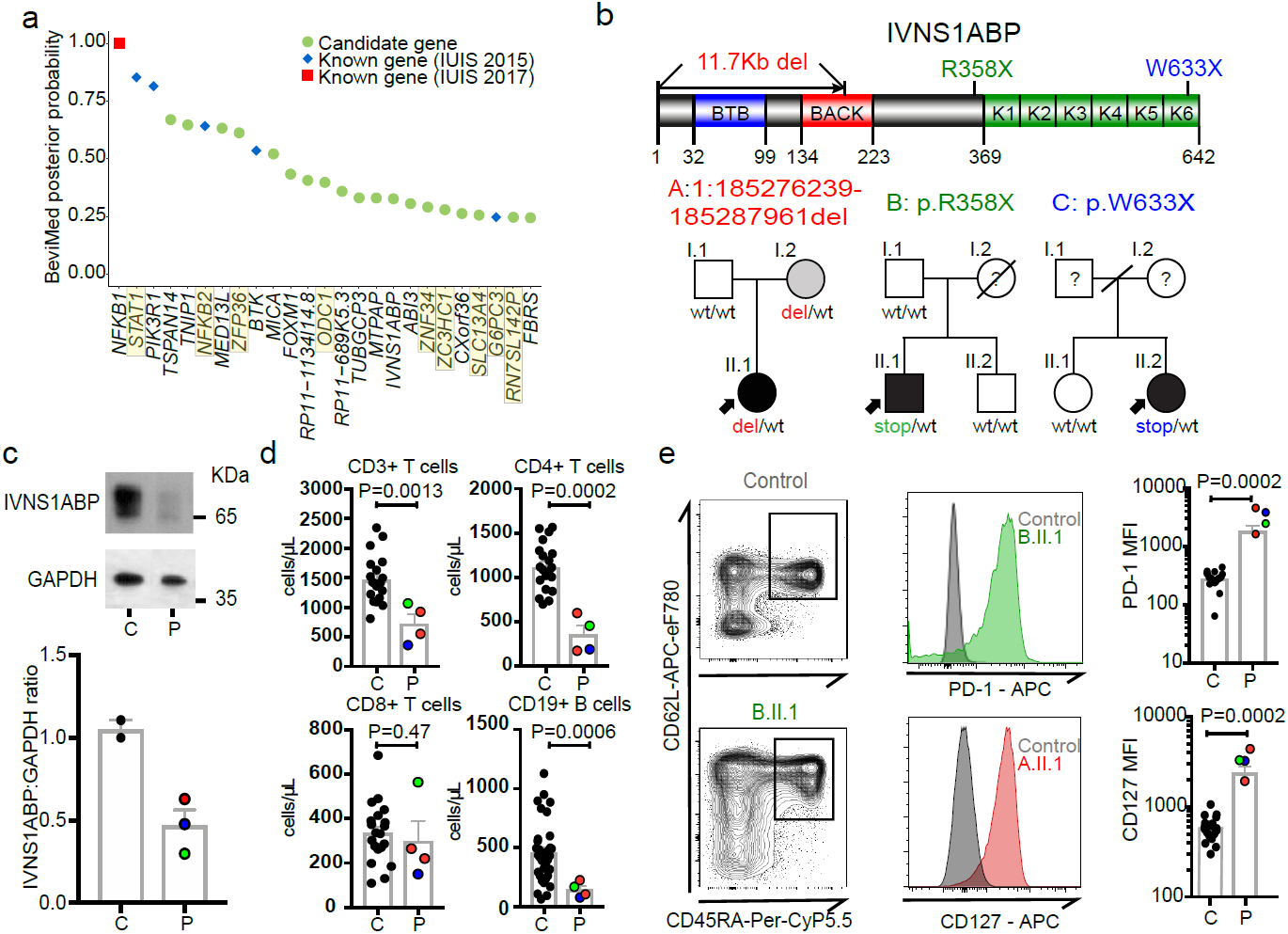
Discovery of novel PID genes in a large cohort WGS analysis. **(a)** BeviMed assessment of enrichment for candidate disease-causing variants in individual genes, in the PID cohort relative to the rest of the NBR-RD cohort (cases n=886, controls n= 9,284). The top 25 candidate genes are shown. Genes highlighted in yellow are those flagged as potentially confounded by population stratification (see **Supplementary Note 2**). Prioritized genes known to cause PID according to the International Union of Immunological Societies (IUIS) in 2015 (blue)^10^ and 2017 (red)^3^. **(b)** Pedigrees of 3 unrelated kindreds with damaging *IVNS1ABP* variants and linear protein position of variants. **(c)** Western blot of IVNS1ABP and GAPDH in whole cell lysates of PBMCs. (Top) Representative blot from A.II.1 (P) and Control (C). For gel source data, see Supplementary Figure 1. (Bottom) Graph of relative IVNS1ABP normalized to GAPDH. (representative of 4 independent experiments). **(d)** Immunophenotyping of CD3+ T cells, CD4+, CD8+ T cells, and CD19+ B cells in C = healthy controls (n=20) and P = *IVNS1ABP* patients (n=4). **(e)** Assessment of CD127 and PD-1 expression in naïve T cells. (Left) Representative gating of naïve (CD45RA+ CD62L+) CD4+ T cells in a control and B.II.1.(Middle) FACS histograms of PD-1 and CD127 from controls and IVNS1ABP patients (B.II.1 and A.II.1). (Right) PD-1 and CD127 mean fluorescence intensity (MFI) values from controls (C, n=20) and patients (P, n=4). All tests two-sided Mann Whitney U. Lines present means, bars = S.E.M.

This method produces a *posterior probability* of association, therefore it is inevitable that, where this is <1, some genes identified will not end up being found to be causal. Such false positives are an integral feature of a method which does not provide statistical proof of causality, but rather ranks/prioritises genes for subsequent functional assessment. They can be minimised by ensuring reasonable assumptions in the Bayesian algorithm^4^, and by taking care to detect and minimise relatedness and population stratification (detailed in **Methods, Supplementary Note 2** and **Supplementary Table 2**).

*NFKB1* and *ARPC1B* were first associated with PID in the literature as a result of familial co-segregation studies^17,18^, and were highly ranked in the BeviMed analysis, validating it as a gene-discovery tool in PID. *NFKB1* had the strongest probability of association (PPA=1-(1.25×10^−8^)), driven by truncating heterozygous variants in 13 patients – leading to our report of *NFKB1* haploinsufficiency as the commonest monogenic cause of CVID^19^. Association of *ARPC1B* with PID (PPA=0.18) was identified by BeviMed based on two recessive cases; one the first reported to link this gene to PID^18^ and the other described below.

To further demonstrate the effectiveness of BeviMed at prioritizing PID-related genetic variants in the cohort, we selected *IVNS1ABP* for validation. BeviMed enrichment (PPA=0.33) of *IVNS1ABP* was driven by three independent heterozygous protein-truncating variants, suggesting haploinsufficiency, while no such variants were observed in controls (**Fig. 2b**). A pathogenic role for *IVNS1ABP* was supported by its intolerance to loss-of-function (pLI=0.994) and a distinctive clinical similarity between the patients – all had severe warts (**Supplementary Note 1**). IVNS1ABP protein expression was around 50% of control, consistent with haploinsufficiency (**Fig. 2c**). The patients also shared a previously undescribed peripheral leukocyte phenotype – with low/normal CD4+ T cells and B cells and aberrant increased expression of CD127 and PD-1 on naïve T cells **(Fig. 2d**,**e)**. Taken together, these data implicate *IVNS1ABP* haploinsufficiency as a novel monogenic cause of PID (**Supplementary Note 1**).

The identification of both known and new PID genes using BeviMed underlines its effectiveness in cohorts of unrelated patients with sporadic disease. As the PID cohort grows, even very rare causes of PID should be detectable with a high positive predictive value (**Extended Data Fig. 3**).

## Identification of regulatory elements contributing to PID

Sequence variation within non-coding regions of the genome can have profound effects on gene expression and would be expected to contribute to susceptibility to PID. We combined rare variant and large deletion (>50bp) events with a tissue-specific catalogue of cis-regulatory elements (CREs)^20^, generated using promoter capture Hi-C (pcHi-C)^21^, to prioritise putative causal PID genes (**Methods**). We limited our initial analysis to rare large deletions overlapping exon, promoter or ‘super-enhancer’ CREs of known PID genes. No homozygous deletions affecting CREs were identified, so we sought individuals with two or more heterozygous variants comprising a CRE deletion with either a rare coding variant or another large deletion in a pcHi-C linked gene. Such candidate compound heterozygote (cHET) variants had the potential to cause recessive disease. Out of 22,296 candidate cHET deletion events, after filtering by MAF, functional score and known PID gene status, we obtained 10 events (**Supplementary Table 3, Extended Data Fig. 4**); the confirmation of three is described.

The *LRBA* and *DOCK8* cHET variants were functionally validated (**Extended Data Figs. 4** and **5**). In these two cases SV deletions encompassed both non-coding CREs and coding exons, but the use of WGS PID cohorts to detect a contribution of CREs confined to the non-coding genome would represent a major advance in PID pathogenesis and diagnosis. *ARPC1B* fulfilled this criterion, with its BeviMed association partially driven by a patient cHET for a novel p.Leu247Glyfs*25 variant resulting in a premature stop, and a 9Kb deletion spanning the promoter region including an untranslated first exon (**Fig. 3a**) that has no coverage in the ExAC database (http://exac.broadinstitute.org). Two unaffected first-degree relatives were heterozygous for the frameshift variant, and two for the promoter deletion (**Fig. 3b**), confirming compound heterozygosity in the patient. Western blotting demonstrated complete absence of ARPC1B and raised ARPC1A in platelets^22^(**Fig. 3c**). *ARPC1B* mRNA was almost absent from mononuclear cells in the patient and was reduced in a clinically unaffected sister carrying the frameshift mutation (**Supplementary Note 1**). An allele specific expression assay demonstrated that the promoter deletion essentially abolished mRNA expression (**Supplementary Note 1**). ARPC1B is part of the Arp2/3 complex necessary for normal actin assembly in immune cells^23^, and monocyte-derived macrophages from the patient had an absence of podosomes, phenocopying deficiency of the Arp2/3 regulator WASp (**Fig. 3d**).

**Figure 3.**
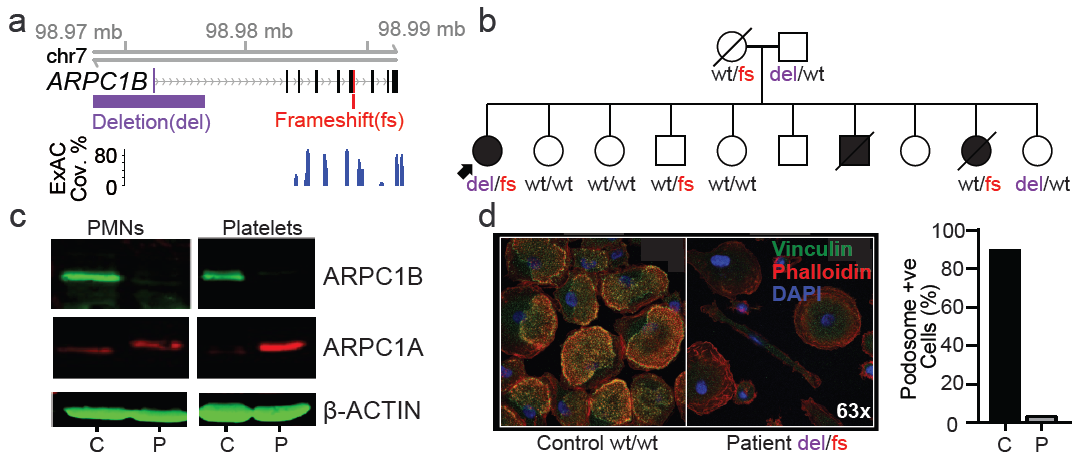
Assessment of WGS data for regulatory region deletions that impact upon PID. **(a)** Genomic configuration of the *ARPC1B* gene locus highlighting the compound heterozygous gene variants. ExAC shows that the non-coding deletion is outside of the exome-targeted regions. **(b)** Pedigree of patient in (a) and co-segregation of *ARPC1B* genotype (wt – wild-type, del – deletion, fs – frameshift). **(c)** Western blot of ARPC1A and ARPC1B in neutrophil and platelet lysates from the patient (P) and control (C, n=1). For gel source data, see Supplementary Figure 1. **(d)** Podosomes were identified by staining adherent, fixed monocyte-derived macrophages for vinculin, phalloidin and the nuclear stain DAPI. Quantification was performed by counting podosomes on at least 100 cells per sample from 10 fields of view at 60x magnification.

While examples of bi-allelic coding variants have been described as causing PID (e.g.^24,25^), here we demonstrate the utility of WGS for detecting compound heterozygosity for a coding variant and a non-coding CRE deletion - a further advantage of a WGS approach to PID diagnosis. Improvements in analysis methodology, cohort size and better annotation of regulatory regions will be required to explore the non-coding genome more fully and discover further disease-causing genetic variants.

## GWAS of the WGS cohort reveals PID-associated loci

The diverse clinical phenotype and variable within-family disease penetrance of PID may be in part due to stochastic events (e.g. unpredictable pathogen transmission) but may also have a genetic basis. We therefore performed a GWAS of common SNPs (minor allele frequency (MAF)>0.05), restricted to 733 AD-PID cases (**Fig. 1a**) to reduce phenotypic heterogeneity (see **Methods**), and 9,225 unrelated NBR-RD controls, and performed a fixed effect meta-analysis of this AD-PID GWAS with a previous CVID study ImmunoChip study (778 cases, 10,999 controls)^8^. This strengthened known MHC and 16p13.13 associations^8^, and found suggestive associations including at 3p24.1 within the promoter region of *EOMES* and at 18p11.21 proximal to *PTPN2*. We also examined SNPs of intermediate frequency (0.005<MAF<0.05) in AD-PID, identifying *TNFRSF13B* p.Cys104Arg variant^26^ (OR=4.04, P = 1.37×10^−12^) (**Fig. 4a, Extended Data Table 3, Extended Data Fig. 6, Supplementary Note 3**). Conditional analysis of the MHC locus revealed independent signals at the Class I and Class II regions, driven by amino-acid changes in the *HLA-B* and HLA-*DRB1* genes known to impact upon peptide binding (**Extended Data Fig. 7**). We next examined the enrichment of non-MHC AD-PID associations in 9 other diseases, finding enrichment for allergic and immune-mediated diseases (IMD), suggesting that dysregulation of common pathways contributes to susceptibility to both (**Supplementary Note 4**).

**Figure 4.**
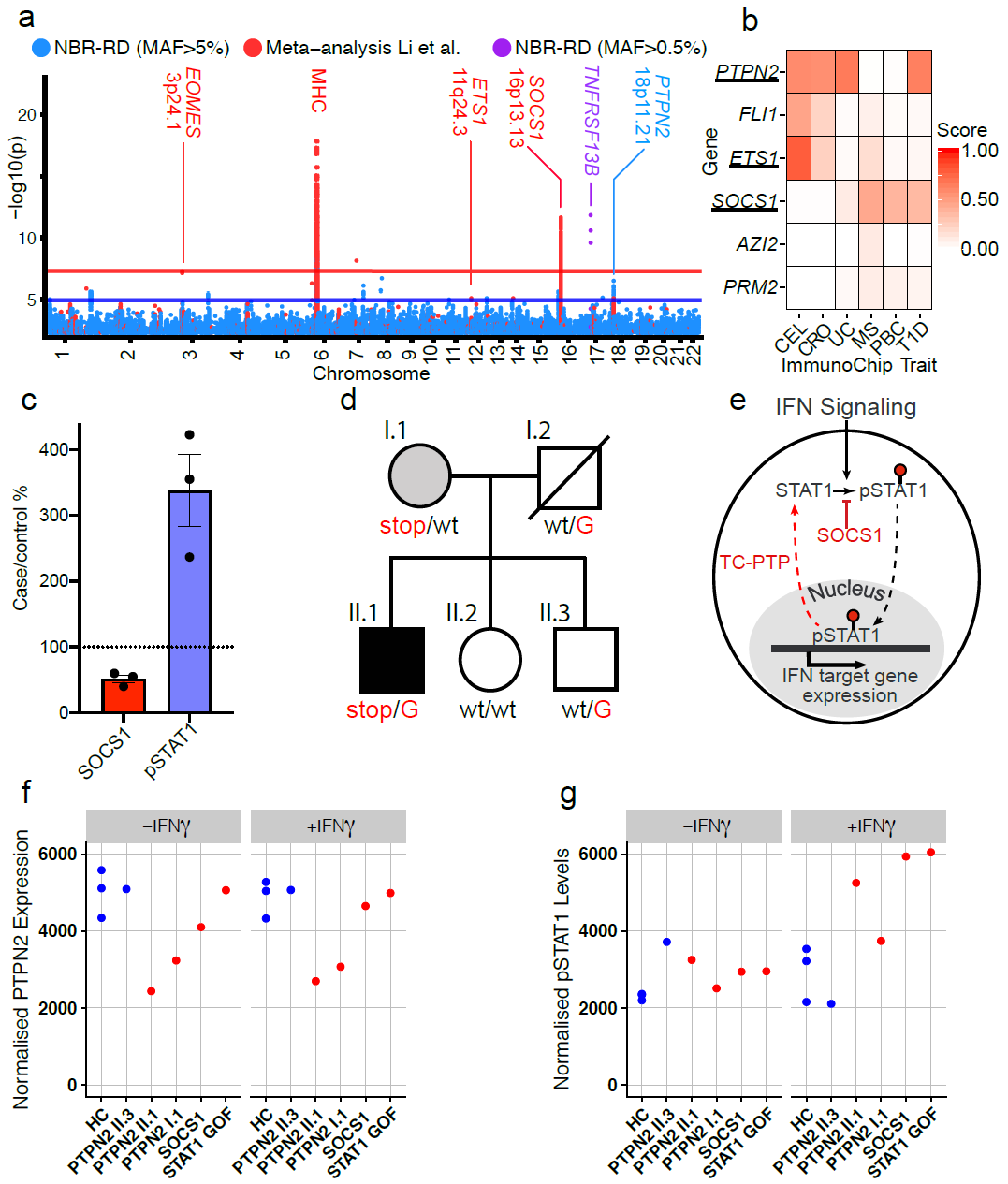
Antibody deficiency (AD-PID) GWAS identifies common variants that mediate disease risk and suggests novel monogenic candidate genes. **(a)** A composite Manhattan plot for the AD-PID GWAS. Blue – common variants (MAF>0.05) analysed in this study (NBR-RD) only (cases n=773, controls n=9,225), red – variants from fixed effects meta-analysis with data from Li *et al*. (cases n=1,511, controls n=20,224); and purple – genome-wide significant low frequency (0.005<MAF<0.05) variants in *TNFRSF13B* locus. Loci of interest are labelled with putative causal protein coding gene names. **(b)** COGS prioritisation scores of candidate monogenic causes of PID using previous autoimmune targeted genotyping studies (**Supplementary Table 4**) across suggestive AD-PID loci (n=4). For clarity, only diseases prioritising one or more genes are shown. CEL – coeliac disease, CRO-Crohn’s disease, UC – ulcerative colitis, MS – multiple sclerosis, PBC – primary biliary cirrhosis and T1D – type 1 diabetes **(c)** Graph of relative pSTAT1 and SOCS1 in lysates made from 2 hour IFN-γ treated T cell blasts from SOCS1 mutation patients and controls. (Lines present mean, error bars=S.E.M.) **(d)** The pedigree of the *PTPN2* mutation patient. Carriers of the rs2847297-G risk allele are indicated. **(e)** Simplified model of how SOCS1 and TC-PTP limit the phosphorylated-STAT1 triggered by interferon signalling. **(f)** Graph of relative PTPN2 and pSTAT1 from the indicated patients and controls, in lysates made from T cell blasts incubated ± IFN-γ for 2 hours. (PTPN2 normalized to tubulin level, pSTAT1 normalised to STAT1 levels, representative of 2 independent experiments)

## GWAS data allows identification of candidate monogenic PID genes and disease-modifying variants

To investigate whether loci identified by GWAS of AD-PID and other IMD might be used to prioritize novel candidate monogenic PID genes, we used the data-driven pcHiC omnibus gene score (COGS) approach^21^ (**Methods, Supplementary Table 4**). We selected six protein-coding genes with above average prioritisation scores in one or more diseases (**Fig. 4b**), and identified a single protein truncating variant in each of *ETS1, SOCS1* and *PTPN2* genes, all occurring exclusively in PID patients. *SOCS1* and *PTPN2* variants were analysed further.

*SOCS1* limits phosphorylation of targets including STAT1, and is a key regulator of IFN-γ signalling^27^. The patient with a heterozygous *de-novo* protein-truncating *SOCS1* variant (p.Met161Alafs*46) presented with CVID complicated by lung and liver inflammation. GeneMatcher^28^ identified an independent pedigree with a protein truncating variant p.Tyr64* in *SOCS1*. All patients showed low/normal numbers of B cells, a Th1-skewed memory CD4+ population and reduced T regulatory (Treg) cells (**Supplementary Note 1**). *Socs1* haploinsufficient mice also demonstrate B lymphopenia^27,29^, a Th1 skew, decreased Tregs^30^ and immune-mediated liver inflammation^31^. In patients’ T cell blasts, SOCS1 was reduced and IFN-γ induced STAT1 phosphorylation was increased (**Fig. 4c**). Taken together this is consistent with *SOCS1* haploinsufficiency causing PID. The initial patient also carried the *SOCS1* pcHiC-linked 16p13.13 risk-allele identified in the AD-PID GWAS (**Supplementary Note 3**) in *trans* with the novel *SOCS1*-truncating variant (**Supplementary Note 1**); such compound heterozygosity suggests common and rare variants might combine to impact upon disease phenotype, a possibility explored further below.

A more detailed example of an interplay between rare and common variants is provided by a family containing *PTPN2* variants (**Fig. 4d**). *PTPN2* encodes the non-receptor T-cell protein tyrosine phosphatase (TC-PTP) that negatively regulates immune responses by dephosphorylation of proteins mediating cytokine signalling. *PTPN2* deficient mice are B cell lymphopenic^32,33^ and haematopoietic deletion leads to B and T cell proliferation and autoimmunity^34^. A novel premature stop-gain at p.Glu291 was identified in a “sporadic” case presenting with CVID at age 20; he had B lymphopenia, low IgG, rheumatoid-like polyarthropathy, severe recurrent bacterial infections, splenomegaly and inflammatory lung disease. His mother, also heterozygous for the *PTPN2* truncating variant, had systemic lupus erythematosus (SLE), insulin-dependent diabetes mellitus, hypothyroidism and autoimmune neutropenia (**Supplementary Note 1**). Gain-of-function variants in *STAT1* can present as CVID (**Supplementary Table 1**) and TC-PTP, like SOCS1, reduces phosphorylated STAT1 (**Fig. 4e**). Both mother and son demonstrated reduced T cell TC-PTP expression and STAT1 hyperphosphorylation, more pronounced in the index case and similar to both SOCS1 haploinsufficient and STAT1 GOF patients (**Fig. 4f**). Thus *PTPN2* haploinsufficiency represents a new cause of PID that acts, at least in part, through increased phosphorylation of STAT1. Reports that use of the Janus Kinase 1 and 2 inhibitor ruxolitinib is effective in controlling autoimmunity in *STAT1*-GOF patients^35^, suggests it might be effective in *SOCS1* and *PTPN2* deficiency.

The index case, but not his mother, carried the G allele of variant rs2847297 at the *PTPN2* locus, an expression quantitative trait locus (eQTL)^36^ previously associated with rheumatoid arthritis^37^. His brother, healthy apart from severe allergic nasal polyposis, was heterozygous at rs2847297 and did not inherit the rare variant (**Fig. 4d**). Allele-specific expression analysis demonstrated reduced *PTPN2* transcription from the rs2847297-G allele, explaining the lower expression of TC-PTP and greater persistence of pSTAT1 in the index case compared to his mother (**Fig. 4g**). This could explain the variable disease penetrance in this family, with *PTPN2* haploinsufficiency alone driving autoimmunity in the mother, but the additional impact of the common variant on the index case causing immunodeficiency. The family illustrates the strength of cohort-wide WGS approach to PID diagnosis, by revealing both a new monogenic cause of disease, and how the interplay between common and rare genetic variants may contribute to the variable clinical phenotypes of PID.

In summary, we show that cohort-based WGS in PID is a powerful approach to provide diagnosis of known genetic defects, and discover new coding and non-coding variants associated with disease (comparison of WGS with other methodologies; **Supplementary Note 5)**. Improved analysis methodology and better integration of parallel datasets, such as GWAS and cell surface or metabolic immunophenotyping, will allow further exploration of the non-coding space, enhancing diagnostic yield. Such an approach promises to transform our understanding of genotype-phenotype relationships in PID and related immune-mediated conditions, and could redefine the clinical boundaries of immunodeficiency, add to our understanding of human immunology, and ultimately improve patient outcomes.

## Methods

### PID cohort

The PID patients and their family members were recruited by specialists in clinical immunology across 26 hospitals in the UK, and one each from the Netherlands, France and Germany. The recruitment criteria were intentionally broad, and included the following: clinical diagnosis of common variable immunodeficiency disorder (CVID) according to internationally established criteria (**Extended Data Table 1**); extreme autoimmunity; or recurrent and/or unusual severe infections suggestive of defective innate or cell-mediated immunity. Patients with known secondary immunodeficiencies caused by cancer or HIV infection were excluded. Although screening for more common and obvious genetic causes of PID prior to enrolment into this WGS study was encouraged, it was not a requirement. Consequently, a minority of patients (16%) had some prior genetic testing, from single gene Sanger sequencing or MLPA to a gene panel screen. Paediatric and familial cases were less frequent in our cohort, in part reflecting that genetic testing is more frequently performed in more severe cases: 31% of paediatric onset cases had prior genetic testing compared to 10% of adult index cases (**Extended Data Fig. 2**).

To expedite recruitment a minimal clinical dataset was required for enrolment, though more detail was often provided. There was a large variety in patients’ phenotypes, from simple “chest infections” to complex syndromic features, and the collected phenotypic data of the sequenced individuals ranged from assigned disease category only to detailed clinical synopsis and immunophenotyping data. The clinical subsets used to subdivide PID patients were based on ESID definitions, as shown in **Extended Data Table 1**. The final PID cohort that we sequenced comprised of 886 index cases, 88 affected relatives, and 344 family members unaffected at the time of recruitment.

To facilitate GWAS analysis by grouping patients with a degree of phenotypic coherence while excluding some distinct and very rare clinical subtypes of PID that may have different aetiologies, a group of patients was determined to have antibody deficiency-associated PID (AD-PID). This group comprised 733 of the 886 unrelated index cases, and included all patients with CID, CVID or Antibody Defect ticked on the recruitment form, together with patients requiring IgG replacement therapy and those with specified low levels of IgG/A/M. SCID patients satisfying these AD criteria were not assigned to the AD-PID cohort.

### WGS data processing

Details of DNA sample processing, whole genome sequencing, data processing pipeline, quality checks, alignment and variant calling, ancestry and relatedness estimation, variant normalisation and annotation, large deletion calling and filtering, and allele frequency calculations, are described in^38^. Briefly, DNA or whole blood EDTA samples were processed and quality checked according to standard laboratory practices and shipped on dry ice to the sequencing provider (Illumina Inc, Great Chesterford, UK). Illumina Inc performed further QC array genotyping, before fragmenting the samples to 450bp fragments and processing with the Illumina TruSeq DNA PCR-Free Sample Preparation kit (Illumina Inc., San Diego, CA, USA). Over the three-year duration of the sequencing phase of the project, different instruments and read lengths were used: for each sample, either 100bp reads on three HiSeq2500 lanes; or 125bp reads on two HiSeq2500 lanes; or 150bp reads on a single HiSeq X lane. Each delivered genome had a minimum 15X coverage over at least 95% of the reference autosomes. Illumina performed the alignment to GRCh37 genome build and SNV/InDel calling using their Isaac software, while large deletions were called with their Manta and Canvas algorithms. The WGS data files were received at the University of Cambridge High Performance Computing Service (HPC) for further QC and processing by our Pipeline team.

For each sample, we estimated the sex karyotype and computed pair-wise kinship coefficients (full methods described in ^47^), which allowed us to identify sample swaps and unintended duplicates, assign ethnicities, generate networks of closely related individuals (sometimes undeclared relatives from across different disease domains) and a maximal unrelated sample set (for the purposes of allele frequency estimation and control dataset in case-control analyses). Variants in the gVCF files were normalised and loaded into an HBase database, where Overall Pass Rate (OPR) was computed within each of the three read length batches, and the lowest of these OPR values (minOPR) assigned to each variant. The rare variant analyses presented here are based on SNVs/InDels with minOPR>0.98. Variants were annotated with Sequence Ontology terms according to their predicted consequences, their frequencies in other genomic databases (gnomAD, UK10K, 1000 Genomes), if they have been associated with a disease according to the HGMD Pro database, and internal metrics (AN, AC, AF, OPR).

Large deletions (those >50bp in length, defined by Illumina) were merged and analysed collectively, as described in^38^. Briefly, sample-level calls by the two algorithms, Manta (which uses read and mate-pair alignment information) and Canvas (which relies on read depth and is optimised for calls >1kb in length), were combined according to a set of rules^38^ to generate a high quality set for each sample (and a large number across the project was visually inspected to ensure reasonably high specificity). To exclude common deletions from further rare variant analyses, we included only those that were observed in fewer than 3% of the samples, as described previously^39^.

### Diagnostic reporting

We screened all genes in the International Union of Immunological Societies (IUIS) 2015 classification for previously reported or likely pathogenic variants. SNVs and small InDels were filtered based on the following criteria: OPR>0.95; having a protein-truncating consequence, gnomAD AF<0.001 and internal AF<0.01; or present in the HGMD Pro database as DM variant. Large deletions called by both Canvas and Manta algorithms, passing standard Illumina quality filters, overlapping at least one exon, and classified as rare by the SVH method were included in the analysis. In order to aid variant interpretation and consistency in reporting, phenotypes were translated into Human Phenotype Ontology (HPO) terms as much as possible. Multi-Disciplinary Team (MDT) then reviewed each variant for evidence of pathogenicity and contribution to the phenotype, and classified them according to the American College of Medical Genetics (ACMG) guidelines^11^. Only variants classified as Pathogenic or Likely Pathogenic were systematically reported, but individual rare (gnomAD AF<0.001) or novel missense variants that BeviMed analysis (see below) highlighted as having a posterior probability of pathogenicity >0.2 were additionally considered as Variants of Unknown Significance (VUS). If the MDT decided that they were likely to be pathogenic and contribute to the phenotype, they were also reported (**Supplementary Table 2**). All variants and breakpoints of large deletions reported in this study were confirmed by Sanger sequencing using standard protocols.

### BeviMed

We used BeviMed^4^ to evaluate the evidence for association, in genetically unrelated individuals, between case/control status and rare genetic variants in a locus. For each gene, we inferred a posterior probability of association (PPA) under Mendelian inheritance models (dominant and recessive), and different variant selection criteria (“moderate” and “high” impact variants based on functional consequences predicted by the Variant Effect Predictor^40^). We inferred a PPA across all association models and the mode of inheritance corresponding to the association model with the greatest posterior probability. We used MAF<0.001 and CADD>=10 as these were selection criteria for rare, likely pathogenic variants used in diagnostic reporting. Approximately 1% of all genes (276/31,350^10^) have previously been implicated as monogenic causes of PID, and we therefore assumed that a few hundred genes are causal of PID overall. We encoded this assumption conservatively, by assigning a prior probability of 0.01 to the association model for each gene. In addition, we used the default prior (mean=0.85) on the “penetrance” parameter, which represents disease risk for individuals carrying pathogenic configuration of alleles at a gene locus (see ^4^ for a detailed description of all parameters and their default values). We then gave all four combinations of inheritance model and variant selection criteria equal prior probability of association of 0.0025 (1/4 of 0.01). We used uniform priors to ensure that our results did not depend on any knowledge of previous gene or variant associations with disease. We obtained a BeviMed PPA for 31,350 genes in the human genome; the highest ranked genes are shown in **Fig. 2a, Supplementary Note 2** and **Supplementary Table 2**. Overall, genes with BeviMed PPA>0.1 were strongly enriched for known PID genes (odds ratio = 15.1, P = 3.1×10^−8^ Fisher’s Exact test), demonstrating that a statistical genetic association approach can identify genes causal for PID.

Conditional on the association model with the highest posterior probability, the posterior probability that each rare variant is pathogenic was also computed. We used a variant-level posterior probability of pathogenicity >0.2 to select potentially pathogenic missense variants in known PID genes to report back. As detailed in Greene *et al.* (Figure 1 in ^4^) the method was calibrated as part of a simulation study estimating positive predictive value (1-FDR) given a fixed level of power. We then examined the relationship between BeviMed rank and ‘known’ gene status in the top fifty genes reported; genes with the highest PPA were significantly enriched for known genes (P<0.008 one-sided Wilcoxon rank-sum test). BeviMed’s sensitivity in prioritizing genes as causal, even if variants exist in only a few cases, is demonstrated by the observation that of the 8 IUIS-defined causal PID genes in the top 50 (all with a BeviMed PPA>0.2), 3 are driven by 2 or 3 cases, while 5 have between 4 and 16.

As allele frequency datasets for non-Europeans are much smaller than for Europeans, potential false positives may be induced by the unintentional inclusion of rare variants observed only in non-European populations^41^.Furthermore, whilst the BeviMed analysis was restricted to the set of cases and controls carefully filtered to minimise relatedness, it remains possible that some associations could be false positives due to residual population stratification. We addressed this by flagging variants whose prioritisation was dependent upon cases with non-European ancestry. In addition, where identical ultra-rare variants were shared between cases, we examined the possibility of cryptic relatedness by seeking direct evidence of shared genetic background (**Supplementary Note 2**). These procedures found that population stratification might contribute to the prioritization of 9 candidate genes among the top 25, as highlighted in **Fig. 2a** and **Supplementary Table 2**. Six of these were novel candidates, but that 3 were known causes of PID indicated that population stratification does not always generate false positives – and implicated genes should therefore be flagged rather than excluded from the list. This potential impact of population stratification underlines the importance of subsequent validation of prioritized genes in order to demonstrate causality.

The BeviMed probabilistic model, based on dominant and recessive inheritance involving a mixture of pathogenic and benign variants, differs from other popular frequentist methods such as SKAT, and is well-suited to the rare disease scenario. When trained on our dataset, SKAT and BeviMed both identified *NKFB1* as the gene with the strongest association signal, but BeviMed placed 8 IUIS 2017 PID genes in the top 50 results whilst SKAT placed 5, and *ARPC1B* was ranked 38th by BeviMed and 289th by SKAT (out of a total of 31,350 tested genes), consistent with the superiority of BeviMed over SKAT and related methods demonstrated in Greene *et al.*^1^.

### Immunohistochemistry: podosome analysis

Frozen peripheral blood mononuclear cells (PBMCs) from healthy donors and patients were thawed and CD14^+^ cells selected using magnetic beads (Miltenyi). 2 × 10^5^ cells/ well in a 24 well plate were seeded on 10ug/ml fibronectin-coated cover slips (R&D systems) in 500ul 20ng/ml macrophage colony stimulating factor (MCSF, Gibco) for 6 days to obtain monocyte-derived macrophages (MDMs). Cells were fixed with paraformaldehyde 4% (Thermo Fisher Scientific) for 10 minutes on ice followed by 8% for 20 minutes at room temperature, permeabilised with 0.1% triton (Sigma) for 5 minutes at room temperature and non-specific binding reduced by blocking with 5% BSA/PBS for 1 hour at room temperature. Cells were incubated with primary anti-vinculin antibody (Sigma 1:200) for 1 hour at room temperature, washed twice with PBS and incubated with secondary antibody conjugated to Alexa Fluor 488 (1:500 Life Technologies) and phalloidin-conjugated to Alexa Fluor 633 (1:200 Thermo Fisher Scientific) for one hour at room temperature. Cells were washed twice with PBS and cover slips mounted onto slides using mounting solution with DAPI for nuclear staining (ProLong Diamond Antifade Mountant with DAPI, Life Technologies) overnight. Slides were imaged using Zeiss 710 confocal microscope at 63x magnification and podosome analysis was carried out on at least 100 cells per sample from 10 fields of view.

### Filtering strategy for candidate regulatory compound heterozygotes

Being underpowered^42^ to detect single nucleotide variants affecting CREs, we limited our initial analysis to large deletions overlapping exon, promoter or ‘super-enhancer’ CREs of known PID genes (**Extended Data Fig. 4**). We selected uncommon (<0.03 frequency NIHR-RD BioResource cohort^38^) large deletion events (>50bp), occurring in PID index cases. We intersected these with a catalogue of of cis-regulatory elements linked to protein-coding genes, created by combining ‘super-enhancer’ and promoter (+/- 500bp window around any protein coding gene transcriptional start site) annotations with promoter capture Hi-C data across 17 primary haematopoietic cell types^21^. Finally, we filtered these events so that only those with linked genes, containing a potentially high impact (CADD>20) rare (MAF<0.001) coding variant, within a previously reported pathogenic gene (IUIS 2017), were taken forward. Events in *ARCPC1B, LRBA* and *DOCK8* were functionally validated. The LRBA cHET variants were confirmed to be in trans by sequencing the parents. Functional LRBA deficiency was demonstrated by impaired surface CTLA-4 expression on Treg cells (**Extended Data Fig. 4**). In the absence of the patient’s mother for sequencing, the DOCK8 variants were confirmed to be in trans by nanopore sequencing and phasing of merged long- and short-read data (see below and **Extended Data Fig. 5**). Functional DOCK8 deficiency was confirmed by a typical clinical phenotype (severe immunodeficiency with prominent wart infection), together with characteristic impaired ex-vivo CD8+, but preserved CD4+, T cell proliferation. The need for rapid bone marrow transplantation has precluded further phenotypic analysis of this patient.

### Phasing of *DOCK8* variants

In order to confirm the phase of two variants detected in the *DOCK8* gene of a single individual, chr9:g. 306626-358548del and chr9:463519G>A, long read sequencing was performed using the Oxford Nanopore Technologies PromethION platform. The DNA sample was prepared using the 1D ligation library prep kit (SQK-LSK109), and genomic libraries were sequenced using a R.9.4.1 PromethION flowcell. Raw signal data in FAST5 format was base called using Guppy (v2.3.5) to generate sequences in FASTQ format, which were then aligned against the GRCh37/hg19 human reference genome using minimap2 (v2.2). Average coverage was 14x and median read length was 4,558 ± 4,007. A high quality set of heterozygous genotypes for the sample was created by using only variants from the short read Illumina WGS data with a phred score of <20 (probability of correct genotype > 0.99). Haplotyping was then performed with Whatshap (v0.14.1) by using the long Nanopore reads to bridge across the informative genotypes from the short read data (https://whatshap.readthedocs.io/en/latest/index.html). We obtained a single high confidence haplotype block spanning the large deletion and the rare missense variant and showing that they were in trans (**Extended data Fig. 5**).

### AD-PID GWAS

GWAS was performed both on the whole PID cohort (N cases = 886) and on a subset comprising AD-PID cases (N cases = 733); the results of the AD-PID analysis were less noisy, and had increased power to detect statistical associations despite a reduced sample size (**Extended Data Fig. 6**). We used 9,225 unrelated samples from non-PID NBR-RD cohorts as controls.

Variants selected from a merged VCF file were filtered to include bi-allelic SNPs with overall MAF>=0.05 and minOPR=1 (100% pass rate across all WGS data for over 13,000 NBR participants). We ran PLINK logistic association test under an additive model. We adjusted for read length to guard against technical differences in genotype calls across the samples sequenced using 100bp, 125bp and 150bp reads, as Illumina chemistries changed throughout the duration of the project. We also used sex and first 10 principal components from the ethnicity analysis as covariates, to mitigate against any population stratification effects. After filtering out SNPs with HWE p<10^−6^, we were left with the total of 4,993,945 analysed SNPs. There was minimal genomic inflation of the test statistic (lambda = 1.022), suggesting population substructure and sample relatedness had been appropriately accounted for. Linear mixed model (LMM) analysis, as implemented in the BOLT-LMM package^43^, is an alternative method of association testing correcting for population stratification. It was used to confirm the observed associations (**Extended Data Table 3**). After genomic control correction^44^ the only genome-wide significant (p<5×10^−8^) signal was at the MHC locus, with several suggestive (p<1×10^−5^) signals (**Extended Data Fig. 6**). We repeated the analysis with more relaxed SNP filtering criteria using 0.005 < MAF < 0.05 and minOPR>0.95 (**Extended Data Fig. 6**). The only additional signal identified were the three *TNFRSF13B* variants shown in **Supplementary Note 3.**

We obtained summary statistics data from the Li et al. CVID Immunochip case-control study^8^ and, after further genomic control correction (lambda = 1.039), performed a fixed effects meta-analysis on 95,417 variants shared with our AD-PID GWAS. Genome-wide significant (p<5×10^−8^) signals were seen at the MHC and 16p13.13 loci, with several suggestive (p<1×10^−5^) signals (**Extended Data Table 3**). After meta-analysis, we conditioned on the lead SNP in each of the genome-wide and suggestive loci by including it as an additional covariate in the logistic regression model in PLINK, to determine if the signal was driven by single or multiple hits at those loci. The only suggestion of multiple independent signals was at the MHC locus (**Extended Data Fig. 7**).

### MHC locus analyses

We imputed classical HLA alleles using the method implemented in the SNP2HLA v1.0.3 package^45^, which uses Beagle v3.0.4 for imputation and the HapMap CEU reference panel. We imputed allele dosages and best-guess genotypes of 2-digit and 4-digit classical HLA alleles, as well as amino acids of the MHC locus genes *HLA-A, HLA-B, HLA-C, HLA-DRB1, HLA-DQA1* and *HLA-DQB1*. We tested the association of both allele dosages and genotypes using the logistic regression implemented in PLINK, and obtained similar results. We then used the best-guess genotypes to perform the conditional analysis (see above), since conditioning is not implemented in PLINK in a model with allele dosages. We repeated the conditional analyses as described above. The results of the sequential conditioning on the two lead classical alleles and amino acids within the Class I and Class II regions are shown in **Extended Data Fig. 7**.

### Allele Specific Expression

RNA and gDNA were extracted from PBMCs using the AllPrep kit (Qiagen) as per the manufacturer’s instructions. RNA was reverse transcribed to make cDNA using the SuperScript™ VILO™cDNA synthesis kit with appropriate minus reverse transcriptase controls, as per the manufacturer’s instructions. The region of interest in the gDNA and 1:10 diluted cDNA was amplified using Phusion (Thermo Fisher) and the following primers on a G-Storm thermal cycler with 30 seconds at 98°C then 35 cycles of 98°C 10 seconds, 60°C 30 seconds, 72°C 15 seconds.

#### ARPC1B

The region of interest spanning the frameshift variant was amplified using the following primers: Forward: GGGTACATGGCGTCTGTTTC / Reverse: CACCAGGCTGTTGTCTGTGA

PCR products were run on a 3.5% agarose gel. Bands were cut out and product extracted using the QIA Quick Gel Extraction Kit (Qiagen), as per protocol. Expected products were confirmed by Sanger sequencing. 4ul fresh PCR product was used in a TOPO^®^cloning reaction (Invitrogen) and used to transform One Shot™ TOP10 chemically competent E. coli. These were cultured overnight then spread on LB agar plates. Individual colonies were picked and genotyped. ARPC1B mRNA expression was assessed using a Taqman gene expression assay with 18S and EEF1A1 as control genes. Each sample was run in triplicate for each gene with a no template control. PCR was run on a LightCycler^®^ (Roche) with 2 mins 50°C, 20 seconds 95°C then 45 cycles of 95°C 3 seconds, 60°C 30 seconds.

#### PTPN2

PTPN2 ASE protocol is modified from above. RNA and genomic DNA were extracted from PBMCs using the AllPrep Kit (Qiagen). RNA was treated with Turbo DNAse (Thermo) and reverse transcribed to generate cDNA using the SuperScript IV VILO master mix (Thermo). The intronic region of interest in gDNA and cDNA was amplified by two nested PCR reactions using Phusion enzyme (Thermo). The primers (F1/R1) and nested primers (F2/R2) used were:

Forward_1: aaagtctggagcaggcagag / Reverse_1: tgggggaactggttatgctttc

Forward_2: ggagctatgatcacgccacatg / Reverse_2: atgctttctggttgggctgac

PCR products were run on a 1% agarose gel. Bands were cut out and product extracted using the QIA Quick Gel Extraction Kit (Qiagen), as per protocol. Expected products were confirmed by Sanger sequencing. 5ng fresh PCR product was used in a TOPO®cloning reaction (Invitrogen) and used to transform One Shot™ TOP10 chemically competent E. coli. These were cultured overnight then spread on LB agar plates. Individual colonies were picked and genotyped. PTPN2 mRNA expression was assessed using a Taqman SNP genotyping assay and on a LightCycler (Roche).

### PAGE and Western Blot analysis

Samples were separated by SDS polyacrylamide gel electrophoresis and transferred onto a nitrocellulose membrane. Individual proteins were detected with antibodies p-STAT1, against STAT1, against SOCS1, against PTPN2 (Cell Signaling Technology, Inc. 3 Trask Lane, Danvers, MA 01923, USA), against ARPC1b (goat polyclonal antibodies, ThermoScientific, Rockford, IL, USA), against ARPC1a (rabbit polyclonal antibodies, Sigma, St Louis, USA) and against actin (mouse monoclonal antibody, Sigma). Secondary antibodies were either donkey-anti-goat-IgG IRDye 800CW, Goat-anti-mouse-IgG IRDye 800CW or Donkey-anti-rabbit-IgG IRDye 680CW (LI-COR Biosciences, Lincoln, NE, USA). Quantification of bound antibodies was performed on an Odyssey Infrared Imaging system (LI-COR Biosciences, Lincoln, NE, USA). Specifically, for IVNS1ABP, whole cell lysates of peripheral blood mononuclear cells were lysed on ice with LDS NuPAGE (Invitrogen) at a concentration of 10^5^ cells per 15ul of LDS. Lysates were denatured at 70°C for 10 minutes then cooled. Lysates were loaded run on Bis-Tris 4-12% Protein Gels (Invitrogen) then transferred to a PVDF membrane (Invitrogen) using iBlot 2 Dry Blotting System (Thermo Fisher Scientific). Membranes were blocked with 5% milk in 5% tris-buffered saline with 0.01% Tween-20 (TBST) for 1 hour at room temperature then incubated overnight with the primary antibodies anti-GAPDH (Cell Signaling Technology) and anti-IVNS1ABP (Atlas Antibodies). Membranes were then washed 3x with TBST at room temperature then incubated with secondary anti-rabbit HRP-conjugated antibody (Cell Signaling Technology) for 1 hour. Membranes were then washed 3x with TBST and 1x with phosphate buffered saline. Membranes were then exposed with Pierce ECL Western Blotting Substrate (Thermo Fischer Scientific) and developed with CL-XPosure Film (Thermo Fischer Scientific).

### Flow cytometry

Peripheral blood mononuclear cells were prepared for analysis by density centrifugation using Histopaque-1077 (Sigma-Aldrich). The following antibodies were used for flow cytometry immunophenotyping: CD3 – BV605 (Biolegend, San Diego, CA, USA), CD4 – APC-eFluor780 (eBioscience, San Diego, CA,USA), CD8 – BV650 (eBioscience, San Diego, CA,USA), CD25 – PE (eBioscience, San Diego, CA,USA), CD127 – APC (eBioscience, San Diego, CA,USA), CD45RA – PerCP-Cy5.5(eBioscience, San Diego, CA,USA, CD19 – BV450 (BD Bioscience, Franklin Lakes, NJ, USA), CD27 – PE-Cy7 (eBioscience, San Diego, CA,USA, CD62L – APC-eF780 (eBioscience, San Diego, CA,USA, CXCR3 – FITC (Biolegend, San Diego, CA, USA), CXCR5 – AF488 (Biolegend, San Diego, CA, USA), CCR7 – PE (Biolegend, San Diego, CA, USA), PD-1 – APC (eBioscience, San Diego, CA,USA), HLA-DR-eFluor450 (eBioscience, San Diego, CA, USA), IgD – FITC (BD Bioscience, Franklin Lakes, NJ, USA). Flow cytometry analysis was performed on a BD LSRFortessa (BD Bioscience) with FACS Diva software (BD Bioscience) for acquisition, then analysis was performed with FlowJo software (LLC).

### AD-PID GWAS Enrichment

Due to the size of the AD-PID cohort, we were unable to use LD-score regression^46^ to assess genetic correlation between distinct and related traits. We therefore adapted the previous enrichment method ‘blockshifter’^47^ in order to assess evidence for the enrichment of AD-PID association signals in a compendium of 9 GWAS European Ancestry summary statistics was assembled from publicly available data. We removed the MHC region from all downstream analysis [GRCh37 chr6:25-45Mb]. To adjust for linkage disequilibrium (LD), we split the genome into 1cM recombination blocks based on HapMap recombination frequencies ^48^. For a given GWAS trait, for *n* variants within LD block *b* we used Wakefield’s synthesis of asymptotic Bayes factors (aBF)^49^ to compute the posterior probability that the *i*^*th*^ variant is causal (*PPCV*_*i*_) under single causal variant assumptions^50^:

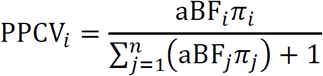

Here *π*_*i*_ = *π*_*j*_are flat prior probabilities for a randomly selected variant from the genome to be causal and we use the value 1×10^−4 51^. We sum over these PPCV within an LD block, *b* to obtain the posterior probability that *b* contains a single causal variant (PPCB).

To compute enrichment for trait *t*, we convert PPCBs into a binary label by applying a threshold such that *PPCB*_*t*_ > 0.95. We apply these block labels for trait *t*, to PPCBs (computed as described above) for our AD-PID cohort GWAS, using them to compute a non-parametric Wilcoxon rank sum statistic, W representing the enrichment. Whilst the aBF approach naturally adjusts for LD within a block, residual LD between blocks may exist. In order to adjust for this and other confounders (e.g. block size) we use a circularised permutation technique^52^ to compute W_null_. To do this, for a given chromosome, we select recombination blocks, and circularise such that beginning of the first block adjoins the end of the last. Permutation proceeds by rotating the block labels, but maintaining AD-PID PPCB assignment. In this way many permutations of W_null_ can be computed whilst conserving the overall block structure.

For each trait we used 10^4^ permutations to compute adjusted Wilcoxon rank sum scores using *wgsea* [https://github.com/chr1swallace/wgsea] R package. For detailed method description see **Supplementary Note 4**.

### PID monogenic candidate gene prioritisation

We hypothesised, given the genetic overlap with antibody associated PID, that common regulatory variation, elucidated through association studies of immune-mediated disease, might prioritise genes harbouring damaging LOF variants underlying PID. Firstly, using summary statistics from our combined fixed effect meta-analysis of AD-PID, we compiled a list of densely genotyped ImmunoChip regions containing one or more variant where P<1×10^−5^. Next, we downloaded ImmunoChip (IC) summary statistics from ImmunoBase (accessed 30/07/2018) for all 11 available studies. For each study we intersected PID suggestive regions, and used COGS (https://github.com/ollyburren/rCOGS) in conjunction with promoter-capture Hi-C datasets for 17 primary cell lines^21,47^ in order to prioritise genes. We filtered by COGS score to select protein coding genes with a COGS score > 0.5, obtaining a list of 11 protein coding genes out of a total of 54 considered.

We further hypothesised that genes harbouring rare LOF variation causal for PID would be intolerant to variation. We thus downloaded pLI scores^53^ and took the product between these and the COGS scores to compute an ‘overall’ prioritisation score across each trait and gene combination. We applied a final filter taking forward only those genes having an above average ‘overall’ score to obtain a final list of 6 candidate genes (Fig. 4d). Finally, we filtered the cohort for damaging rare (gnomAD AF<0.001) protein-truncating variants (frameshift, splice-site, nonsense) within these genes in order to identify individuals for functional follow up.

### Statistical analyses

Statistical analyses were carried out using R (v3.3.3 – “Another Canoe”) and Graphpad Prism (v7) unless otherwise stated. All common statistical tests are two-sided unless otherwise stated. No statistical methods were used to pre-determine sample size

## Acknowledgements

Funding for the NIHR-BioResource was provided by the National Institute for Health Research (NIHR, grant number RG65966). We gratefully acknowledge the participation of all NIHR BioResource volunteers, and thank the NIHR BioResource centre and staff for their contribution. JEDT is supported by the MRC (RG95376 and MR/L006197/1). AJT is supported by the Wellcome Trust (104807/Z/14/Z) and the NIHR Biomedical Research Centre at Great Ormond Street Hospital for Children NHS Foundation Trust and University College London. KGCS is supported by the Medical Research Council (program grant MR/L019027) and is a Wellcome Investigator. AJC was supported by the Wellcome [091157/Z/10/Z], [107212/Z/15/Z], [100140/Z/12/Z], [203141/Z/16/Z]; JDRF [9-2011-253], [5-SRA-2015-130-A-N]; NIHR Oxford Biomedical Research Centre and the NIHR Cambridge Biomedical Research Centre. EE has received funding from the European Union Seventh Framework Programme (FP7-PEOPLE-2013-COFUND) under grant agreement no 609020-Scientia Fellows. ER is supported supported by the Wellcome Trust [201250/Z/16/Z]. DE is supported by the German Federal Ministry of Education and Research (BMBF) within the framework of the e:Med research and funding concept (SysInflame grant 01ZX1306A; GB-XMAP grant 01ZX1709) and funded by the Deutsche Forschungsgemeinschaft (DFG, German Research Foundation) under Germany’s Excellence Strategy – EXC 2167-390884018.

## Author Contributions

JEDT, ES, JS, ZZ, WR, NSG, PT, ER, AJC carried out experiments. HLA, OSB, JEDT, JHRF, DG, IS, CP, SVVD, ASJ, JM, JS, PAL, AGL, KM, EE, DE, SFJ, THK, ET performed computational analysis of the data. HLA, IS, CP, MB, CrS, RL, PJRM, JS, KES conducted sample and data processing. JEDT, ES, WR, MJT, RBS, PG, HEB, AW, SH, RL, MSB, KCG, DSK, AC, DE, AH, NC, SG, AH, SG, SJ, CaS, FB, SS, SOB, TWK, WHO, AJT recruited patients, provided clinical phenotype data and confirmed genetic diagnosis. All authors contributed to the analysis of the presented results. KGCS, JEDT, HLA, WR and OSB wrote the paper with input from all other authors. KGCS, WHO, AJT and TWK conceived and oversaw the research programme.

## Author Information

<u>Members of the NBR-RD PID Consortium:</u> Zoe Adhya, Hana Alachkar, Carl E Allen, Ariharan Anantharachagan, Richard Antrobus, Gururaj Arumugakani, Chiara Bacchelli, Helen E Baxendale, Claire Bethune, Shahnaz Bibi, Barbara Boardman, Claire Booth, Matthew Brown, Michael J Browning, Mary Brownlie, Matthew S Buckland, Siobhan O Burns, Oliver S Burren, Anita Chandra, Ivan K. Chinn, Hayley Clifford, Nichola Cooper, Godelieve J de Bree, E Graham Davies, Sarah Deacock, John Dempster, Lisa A Devlin, Elizabeth Drewe, J David M Edgar, William Egner, Shuayb El Khalifa, Tariq El-Shanawany, James H R Farmery, H Bobby Gaspar, Rohit Ghurye, Kimberly C Gilmour, Sarah Goddard, Pavels Gordins, Sofia Grigoriadou, Scott J Hackett, Rosie Hague, Lorraine Harper, Grant Hayman, Archana Herwadkar, Stephen Hughes, Aarnoud P Huissoon, Stephen Jolles, Julie Jones, Yousuf M Karim, Peter Kelleher, Sorena Kiani, Nigel Klein, Taco W Kuijpers, Dinakantha S Kumararatne, James Laffan, Hana Lango Allen, Sara E Lear, Hilary Longhurst, Lorena E Lorenzo, Paul A Lyons, Jesmeen Maimaris, Ania Manson, Elizabeth M McDermott, Hazel Millar, Anoop Mistry, Valerie Morrisson, Sai H K Murng, Iman Nasir, Sergey Nejentsev, Sadia Noorani, Eric Oksenhendler, Mark J Ponsford, Waseem Qasim, Ellen Quinn, Isabella Quinti, Alex Richter, Crina Samarghitean, Ravishankar B Sargur, Sinisa Savic, Suranjith L Seneviratne, W A Carrock Sewell, Fiona Shackley, Olga Shamardina, Ilenia Simeoni, Kenneth G C Smith, Emily Staples, Hans Stauss, Cathal L Steele, James E Thaventhiran, David C Thomas, Moira J Thomas, Adrian J Thrasher, John A Todd, Anton T J Tool, Salih Tuna, Rafal D Urniaz, Steven B Welch, Lisa Willcocks, Sarita Workman, Austen Worth, Nigel Yeatman, Patrick F K Yong.

Reprints and permissions information is available at www.nature.com/reprints. Readers are welcome to comment on the online version of the paper.

## Ethics Declaration

NBR-RD participants from the UK were consented under the East of England Cambridge South national research ethics committee (REC) reference 13/EE/0325. Participants recruited outside of the UK were consented by the recruiting clinicians under the ethics governance of their respective hospitals.

## Data Availability

WGS and phenotype data from participants is available from one of 3 data repositories determined by the informed consent of the participant. (1) Data from participants enrolled in the NIHR BioResource for the 100,000 Genomes Project–Rare Diseases Pilot can be accessed via Genomics England Limited: https://www.genomicsengland.co.uk/about-gecip/joining-research-community/. (2) data from the UK Biobank samples are available through a data release process overseen by UK Biobank (https://www.ukbiobank.ac.uk/). (3) data from the remaining NIHR BioResource participants is available from the European Genome-phenome Archive (EGA) at the EMBL European Bioinformatics Institute (EGA accession code EGAD00001004523). Patients all fall into group (3) and controls into groups (1)-(3). Variants listed in Supplementary Table 1 (diagnostic findings) have been submitted to ClinVar and are accessible under “NIHR_Bioresource_Rare_Diseases_PID”. Summary statistics are available via GWAS Catalog [Accession number granted upon acceptance of the manuscript].

## Code Availability

R code for running major analyses are available at https://github.com/ollyburren/pid_thaventhiran_et_al.

## Extended Data Figures and Tables

**Extended Data Figure 1 – Graphical abstract**

**Extended Data Figure 2 – Genetic testing in the PID cohort prior to WGS recruitment, in sporadic versus familial cases.** Any type of genetic test is included, such as single exon/gene sequencing, MLPA, or targeted gene panel/exome sequencing. The information was supplied on the referral form and is likely an underestimate of the number of patients with additional genetic testing.

**Extended Data Figure 3 – BeviMed simulation study of Positive Predictive Value (PPV) with increasing disease cohort size.** We simulated genotypes at 25 rare variant sites in a hypothetical locus amongst 20,000 controls and a further 1,000, 2,000, 3,000, 4,000 or 5,000 cases. We simulated that 0.2%, 0.3%, 0.4% or 0.5% of the cases had the hypothetical locus as their causal locus. We distinguish between cases due to the hypothetical locus (CHLs) and cases due to other loci (COLs). The allele frequency of 20 variants was set to 1/10,000 amongst the cases and COLs. The allele frequency of the remaining 5 variants was set to zero amongst the controls and COLs. One of the five variants was assigned a heterozygous genotype amongst the CTLs at random. Thus, we represent a dominant disorder caused by variants with full penetrance. As inference is typically performed across thousands of loci, with only a small number being causal, we assumed a mixture of 100 to 1 non-causal to causal loci. In order to compute the PPV for a given threshold on the posterior probability of association (PPA), we computed PPAs for 10,000 datasets without permutation of the case/control labels and 10,000 further datasets with a permutation of the case/control labels. We then sampled 1,000 PPAs from the permuted set and 10 PPAs from the non-permuted set to compute the PPV obtained when the PP threshold was set to achieve 100% power. The mean over 2,000 repetitions of this procedure is shown on the y-axis. The x-axis shows the number of cases in a hypothetical cohort. As the number of cases increases from 1,000 to 5,000, the PPV increases above 87.5% irrespective of the proportion of cases with the same genetic aetiology. This demonstrates the utility of expanding the size of the PID case collection for detecting even very rare aetiologies resulting in the same broad phenotype as cases with different aetiologies. In practice, the PPV/power relationship may be much better, as the wealth of phenotypic information of the cases can allow subcategorization of cases to better approximate shared genetic aetiologies.

**Extended Data Figure 4 – Candidate cHET filtering strategy and *LRBA* patient. (a)** Filtering strategy to identify candidate compound heterozygous (cHET) pathogenic variants consisting of a rare coding variant in a PID-associated gene and a deletion of a cis-regulatory element for the same gene. **(b)** Regional plot of the compound heterozygous variants. Gene annotations for are taken from Ensembl Version 75, and the transcripts shown are those with mRNA identifiers in RefSeq (ENST00000357115 and ENST00000510413). The position of each variant relative to the gene transcript is shown by a red bar, with the longer bar indicating the extent of the deleted region. Variant coordinates are shown for the GRCh37 genome build. **(c)** Pedigree of LRBA patient demonstrating phase of the causal variants. **(d)** FACS dotplot of CTLA-4 and FoxP3 expression in LRBA cHET patient and a healthy control (representative of 2 independent experiments). Numbers in black are the percentage in each quadrant. Numbers in red are the MFI of CTLA-4 staining in FoxP3 -ve and FoxP3 +ve cells. **(e)** Normalised CTLA-4 expression, assessed as previously described in Hou *et al.* (Blood, 2017), in the LRBA cHET patient (n=1), healthy controls (n=8) and positive control CTLA-4 (n=4) and LRBA (n=3 deficient patients. Horizontal bars indicate mean +/- SEM.

**Extended Data Figure 5 - *DOCK8* cHET patient. (a)** Regional plot of the compound heterozygous variants. Gene annotations for are taken from Ensembl Version 75, and the transcripts shown are those with mRNA identifiers in RefSeq (ENST00000432829 and ENST00000469391). The position of each variant relative to the gene transcript is shown by a red bar, with the longer bar indicating the extent of the deleted region. Variant coordinates are shown for the GRCh37 genome build. **(b)** Photographs of the extensive HPV associated wart infection in the *DOCK8* cHET patient. **(c)** cHET variant phasing. Top: cartoon representation of phasing using high quality heterozygous calls from short read WGS data and long-read nanopore sequencing data. Bottom panel: WGS and nanopore data from the *DOCK8* patient. The two variants (large deletion and missense substitution) are shown in the bottom track (orange), and a single phase block (green) that spans the entire region between the two variants confirmed them to be in-trans. **(d)** Dye-dilution proliferation assessment in response to phytohaemagglutinin (PHA) and anti-CD3/28 beads in CD4+ and CD8+ T cells in patient and control cells (representative of 2 independent experiments). Staining was performed with CFSE dye (Invitrogen, Carlsbad, CA, USA) with the same additional fluorochrome markers as described in the flow cytometry methods section.

**Extended Data Figure 6 – Manhattan plots of (a) all-PID MAF>5%, (b) AD-PID MAF>5% and (c) AD-PID 0.5%<MAF<5% GWAS results.** Sample sizes: all-PID cases n=886; AD-PID cases n=733; controls n=9,225. Each point represents an individual SNP association P-value, adjusted for genomic inflation. Only signals with P<1×10^−2^ are shown. None of the SNPs in plot (c) appear in the results of the common variant GWAS in (b), and are therefore additional signals gained from a GWAS including variants of intermediate MAF. Red and blue lines represent genome-wide (P<5×10^−8^) and suggestive (P<1×10^−5^) associations, respectively. Note the additional genome-wide significant signal representing the *TNFRSF13B* locus, and several suggestive associations that only become apparent with variants in the 0.5% - 5% MAF range shown in (c). Suggestive loci are indicated by the rsID of the lead SNP in each chromosome. Note that lead SNPs in AD-PID GWAS (b) may differ from meta-analysis lead SNPs.

**Extended Data Figure 7 – MHC locus conditional analyses in AD-PID GWAS (cases n=733, controls n=9**,**225). (a)** Locuszoom association plots of AD-PID GWAS MHC locus initial (top) and conditional (middle, bottom) analyses results. The *x* and left *y* axes represent the chromosomal position and the - log10 of the association P-value, respectively. Each point represents an analysed SNP, with the lead SNP indicated by a purple diamond and all other points coloured according to the strength of their LD with the lead SNP. Purple lines represent HapMap CEU population recombination hotspots. The bottom panel shows a selection of genes in the region, with over 150 genes omitted. Top: association plot of the most significant signal rs1265053, which is in the Class I region and close to *HLA-B* and *HLA-C* genes. Middle: plot showing the association remaining upon conditioning on rs1265053, with the strongest signal rs9273841 mapping to the Class II region close to *HLA-DRB1* and *HLA-DQA1* genes. Bottom: plot showing the association signal remaining upon conditioning on both rs1265053 and rs9273841. **(b**,**c)** MHC locus conditional analyses of the classical HLA alleles **(b)** and amino acids of individual HLA genes **(c)**. Each point represents a single imputed classical allele or amino acid, with those marked in red indicating those added as covariates to the logistic regression model: the Class I signal (second row plots), the Class II signal (third row plots), and both Class I and Class II signals (bottom row plots). The HLA allele and amino acid shown in the bottom plots are those with the lowest P-value remaining after conditioning on both Class I and Class II signals; as there are no genome-wide significant signals remaining, the results suggest there are two independent signals at the MHC locus. **(d)** Protein modelling of two independent MHC locus signals: *HLA-DRB1* residue E71 and *HLA-B* residue N114 using PDB 1BX2 and PDB 4QRQ respectively. Protein is depicted in white, highlighted residue in red, and peptide is in green.

**Extended Data Table 1 – ESID definition of PID subtypes**. Participants were defined phenotypically to the groups: primary antibody deficiency, CVID, CID, severe autoimmunity/immune dysregulation, autoinflammatory syndrome, phagocyte disorder, and unspecified PID according to the European Society for Immunodeficiencies (ESID) registry diagnostic criteria (https://esid.org/Working-Parties/Registry-Working-Party/Diagnosis-criteria).

**Extended Data Table 2 – Description of the NIHR BioResource - Primary Immunodeficiency cohort**. High-level clinical description and relevant clinical features were provided by recruiting clinicians. Index cases are patients recruited as sporadic cases or probands in pedigrees, and determined to be genetically unrelated by pairwise comparisons of common SNP genotypes in the WGS data. Numbers in brackets refer to the percentage of index cases in each category. Total number of patients is the sum of index cases and any affected relatives sequenced in this study.

**Extended Data Table 3 – Genome-wide significant (P<5×10**^**-8**^**) and suggestive (P<1×10**^**-5**^**) signals in our AD-PID and Li *et al.* (Nat Comm, 2015) CVID GWAS meta-analysis.** The AD-PID WGS cohort included 733 cases and 9225 controls, whereas the CVID Immunochip cohort included 778 cases and 10999 controls. The total number of shared meta-analysed variants was 95417. P-values are adjusted for individual study genomic inflation factor lambda. The selection of genes from each locus used in COGS analysis is described in Methods and Supplementary Note 3.

