## Supplementary Information for "Whole genome sequencing of a sporadic primary immunodeficiency cohort"

#### for Thaventhiran *et al.*


**Strategy Outline** **2**

**Supplementary Methods** **3**

Enrolment and sample collection

Ethics statement

**Supplementary Notes** **4**

Supplementary Note 1 – Clinical Case Descriptions **4**

a) IVNS1ABP **4**

b) ARPC1B **8**

c) SOCS1 and PTPN2 **10**

Supplementary Note 2 – Population stratification in the PID cohort **14**

Supplementary Note 3 – AD-PID GWAS candidate genes **18**

Supplementary Note 4 – AD-PID GWAS enrichment methods **29**

Supplementary Note 5 – PID diagnosis: a comparison of WGS vs WES+GWAS **34**

**Supplementary Tables – as separate files**

Supplementary Table 1 – Diagnostic PID findings

Supplementary Table 2 – Top BeviMed PID candidate genes

Supplementary Table 3 – Prioritised candidate cHET genes

Supplementary Table 4 – COGS prioritised genes

#### Strategy Outline

This paper addresses the pressing clinical issue that while most Primary Immunodeficiency (PID) patients have sporadic, mild, adult-onset disease associated with B cell abnormalities (such as Common Variable Immunodeficiency, CVID), most genetic diagnoses in PID are made in the smaller subgroup of patients with severe, familial and young-onset disease (such as Severe Combined Immunodeficiency, SCID).

To address this we used whole genome sequencing to study a cohort of PID patients comprised predominantly of the former category, and examined this data using Bayesian methods (BeviMed) and a number of genome-wide approaches. We:

- describe the ability of this cohort-based Bayesian analysis, as an approach distinct from standard familial co-segregation, to identify monogenic causes of sporadic PID;
- demonstrate that integrated cohort-based WGS approaches can begin to find genetic defects in the non-coding genome, from rare and highly penetrant to PID-associated common variants;
- illustrate the potential impact of common variants on the variable penetrance of PID;
- achieve this despite the heterogenous and sometimes poorly-defined clinical phenotypes associated with PID.

### Supplementary Methods

#### Enrolment and sample collection

The main cohort used in this study was recruited and sequenced as part of the NIHR BioResource – Rare Diseases (NBR-RD) project (https://bioresource.nihr.ac.uk/rare-diseases/rare-diseases/), which includes nearly 10,000 patients with rare diseases and their relatives. Primary Immunodeficiency (PID) was one of 15 disease domains under which participants were enrolled between December 2012 and March 2017.

DNA or blood samples were received and processed by the Cambridge Translational GenOmics (CATGO) Laboratory, and those passing the quality checks were sent to Illumina for whole genome sequencing (see Methods for details). Additional WGS data from samples from the Genomics England Ltd (GEL) rare diseases pilot study for the 100,000 Genomes Project (https://www.genomicsengland.co.uk/the-100000-genomes-project/), and from the UK Biobank cohort (http://www.ukbiobank.ac.uk) were processed together with those of the NBR-RD participants, for the purposes of sequence and variant quality checks, overall allele frequency calculations, and as additional controls in statistical analyses.

#### Ethics statement

NBR-RD participants from the UK were consented under the East of England Cambridge South national research ethics committee (REC) reference 13/EE/0325. Participants recruited outside of the UK were consented by the recruiting clinicians under the ethics governance of their respective hospitals.

Samples used in the promoter capture Hi-C experiments were collected with written and signed informed consent. The study was approved by the East of England – Cambridgeshire and Hertfordshire Research Ethics Committee for the project entitled: 'An investigation into genes and mechanisms based on genotype-phenotype correlations in type 1 diabetes and related diseases using peripheral blood mononuclear cells from volunteers that are part of the Cambridge BioResource project' (REC number 05/Q0106/20).

### Supplementary Note 1

### Clinical case descriptions

#### a) *IVNS1ABP* pedigrees and functional assessment

BeviMed analysis identified association of *IVNS1ABP* with PID, driven by three independent cases all with protein-truncating variants (**Figure 1a-c; Table 1**). To investigate these cases further and functionally validate the variants as disease-causing, we first recruited family members and assessed the immune phenotype of the blood cells of the variant-carrying individual and by flow cytometry. This demonstrated all variant carriers had the non-specific finding of low CD4+ T cells and numbers of total T cells and B cells towards the lower limit of the normal range (**manuscript Fig. 2d**). Assessment of T cell differentiation and activation demonstrated no difference between variant carriers and healthy controls (**Figure 2a-e**). However, assessment of the naïve T cells demonstrated increased expression of the CD127 (IL-7 receptor) and aberrant expression of the immune checkpoint regulator PD-1 on CD4+ and CD8+ T cells (**manuscript Fig. 2e; Figure 2f below**) on all *IVNS1ABP* variant carrying individuals. Although PD-1 is expressed on T cells during thymic development[2] its expression in the periphery in unchallenged subjects on naïve T cells has never been reported. This result confirms the full penetrance of IVNS1ABP haploinsufficiency on the immunological phenotype and positions IVNS1ABP as a novel regulator of PD-1 expression.

The clinical histories of the probands of the patients with *IVNS1ABP* BeviMed-identified variants are detailed below. The available relatives of these patients were assessed for the presence of the rare, putative causal, variant (**Figure 1**).

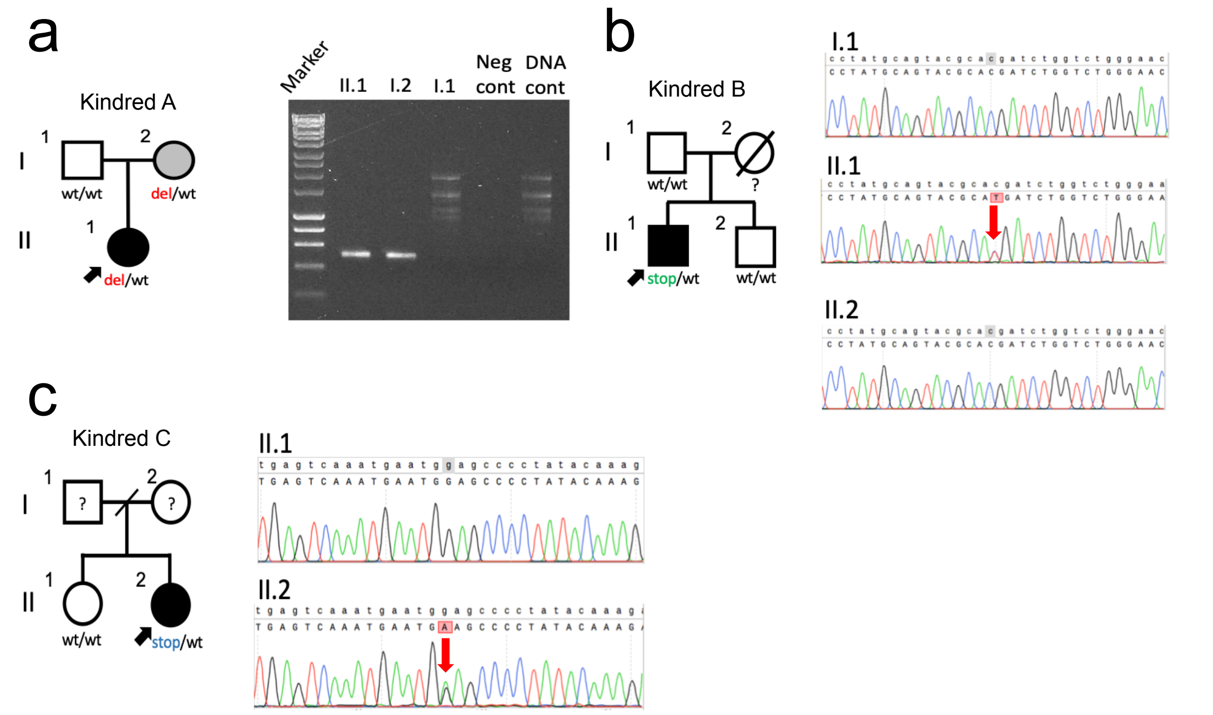


**Figure 1. *IVNS1ABP* pedigrees and Sanger sequences. (a)** Kindred A pedigree and gel image of a single long range PCR experiment targeting the *IVNS1ABP* deletion chr1:g.18276230_185287961 and showing bands representing the deletion only present in index case II.1 and her affected mother I.2. **(b)** pedigree of Kindred B and sanger traces of the p.R358* variant (red arrow). **(c)** Kindred C pedigree and Sanger sequence traces of p.W633* variant (red arrow). ? = unknown genotype and phenotype. Fluorescent traces presented from 1 experiment performed from each family member.

#### Clinical Histories

##### *IVNS1ABP* Kindred A

The proband of Kindred A (A.II.1) presented with bilateral retinal vasculitis at 19-years-old. He had a history of numerous troublesome warts on his face and hands which had been non-responsive to treatments for many years preceding this presentation. On investigation he was found to be CD4+ T cell lymphopenic and HIV testing was negative. Due to a positive interferon-gamma release assay and high-risk history of tuberculous contact he was empirically treated with anti-tuberculous quadruple therapy and prednisolone for presumed tuberculous and bilateral retinitis vasculitis.

Limited clinical information was available from the mother of the proband, the only *IVNS1ABP* variant-carrying family member identified from the pedigree assessments. However, in this individual (kindred A 1.2.) an infection susceptibility was not noted. This could be because of partial penetrance of the variant or because the clinical phenotype is infection susceptibility and a lack of pathogen encounter prevented clinical phenotype expression.

Whole genome sequencing identified a heterozygous 11.7Kb partial gene deletion (chr1:g.185276239_185287961del). The deletion was also present in his mother, who despite reporting no severe infections, did show a low CD4+ T cell and CD19+ B cell count.

##### *IVNS1ABP* Kindred B

Kindred B is of British Caucasian origin. Index case B.II.1 initially presented in childhood with non-infectious diarrhoea. At age 21-years-old he underwent colonoscopy and biopsies showing evidence of an inflammatory colitis. He also suffered with cutaneous boils, few of which required incision and drainage, and warts on the dorsum of his hands. Ha has continued to suffer with recalcitrant warts on his hands and feet, and also had a history also of cutaneous boils, some of which required incision and drainage under anaesthetic. At 30-years-old he suffered 2 episodes of bacterial pneumonia requiring hospitalization and intravenous antibiotics, at which point he was found to be hypogammaglobulinaemic with a low CD4+ T cell count and inverted CD8+ ratio. HIV testing was negative. Chest CT scan revealed basal bronchiectasis attributed to subclinical infections due to hypogammaglobulinaemia. He also had recurrent nasal polyposis which required functional endoscopic sinus surgery on 2 occasions.

Whole genome sequencing identified a rare heterozygous nonsense *IVNS1ABP* variant (chr1:g. 185270152G>A; ENST00000367498 c.1072C>T; p.Arg358*). This variant was not present in his healthy father or brother. His mother had died several years prior to genetic testing from a myocardial infarction.

##### *IVNS1ABP* Kindred C

C.II.2 was referred aged 35-years-old due to longstanding recalcitrant troublesome warts on her hands. The warts were HPV associated and had not responded to numerous therapies, quickly recurring following surgical removal. She had a past medical history of coeliac disease since childhood and achalasia aged 31-years-old. Aged 29-years-old she had had a protracted glandular fever due to primary EBV infection which she attributed to causing long-term fatigue. Her EBV blood viral load has been undetectable on several occasions since resolution of the primary infection. Due to recurrent bacterial sinusitis she had been taking prophylactic doxycycline for several years.

Whole genome sequencing identified a novel heterozygous nonsense *IVNS1ABP* variant (chr1:g. 185267197C>T; ENST00000367498 c.1899G>A; p.Trp633*). Her sister has a history of breast cancer, but no recurrent or severe infections did not carry the variant. Neither of her parents were available for genetic investigation.

| | Age | *IVNS1ABP* variant | Ethnicity | Infections | Other clinical features | (cells/$mm^3$) | | | | | IgG (6.0-16.0g/l) | IgA (0.9-4.5g/l) | IgM (0.5-2.0g/l) |
|---|---|---|---|---|---|---|---|---|---|---|---|---|---|
| | | | | | | LC | CD3+ | CD4+ | CD8+ | CD19+ | | | |
| A.I.2 | 51 | 1:182766239-185287961del | Pakistani | | | 1275 | 923 | 599 | 265 | 108 | | | |
| A.II.1 | 19 | 1:185276239-185287961del | Pakistani | HPV warts | Retinal vasculitis | 1030 | 550 | 174 | 220 | 227 | 11.5 | 2.07 | 0.95 |
| B.II.1 | 56 | c.1072C>T p.R358X | White British | Sinopulmonary bacterial<br>Staphylococcal boils<br>HPV warts | Bronchiectasis<br>Nasal polyposis<br>Inflammatory colitis | 1300 | 1066 | 456 | 564 | 170 | <0.33 (L) | <0.07 (L) | 0.11 (L) |
| C.II.2 | 44 | c.1899G>A p.W633X | White British | EBV<br>HPV warts | Coeliac disease<br>Achalasia<br>Chronic fatigue syndrome<br>Hypothyroidism | 715 | 360 | 190 | 150 | 80 | 11.1 | 0.64 | 1.03 |

**Table 1. Clinical and cellular phenotypes of patients with *IVNS1ABP* variants prioritised by BeviMed.** LC – lymphocytes. (L) – low compared to normal range.

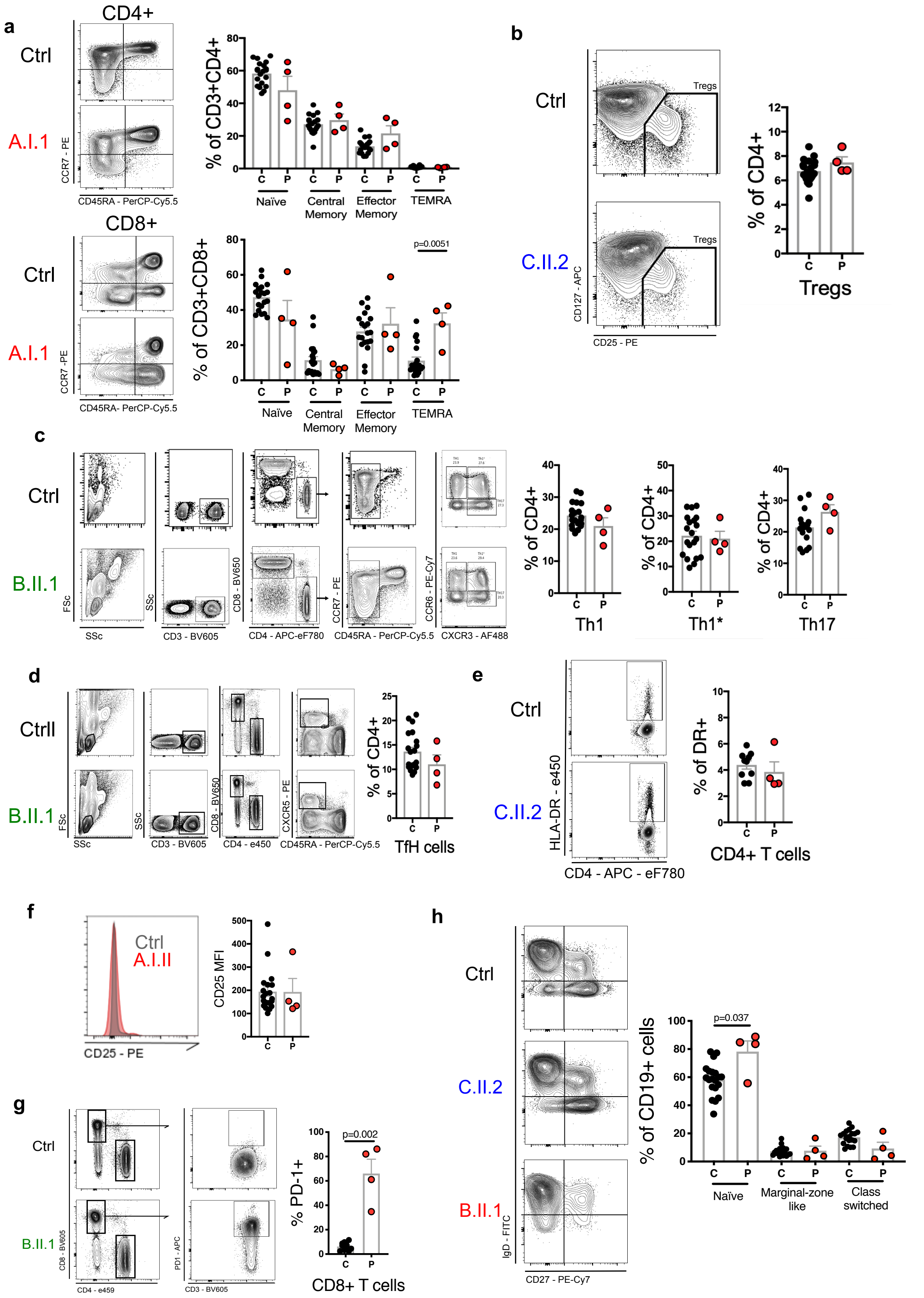


**Figure 2. Extended immunophenotyping of *IVNS1ABP* patients compared with healthy controls. (a)** naïve (CD3+CD4+ or CD8+CD45RA+CCR7+), central memory (CD3+CD4+ or CD8+CD45RA-CCR7+), effector memory (CD3+CD4+ or CD8+CD45RA-CCR7-)and terminal effector memory RA (CD3+CD4+ or CD8+CD45RA+CCR7-) T cells in healthy controls (C) and *IVNS1ABP* patients (P). **(b)** Cells expressing surface markers found on regulatory T cells (Tregs) (CD3+CD4+CD25+CD127$^{low}$) in healthy controls (C) and patients (P). **(c)** Gating strategy for healthy

control and patient for surface markers that characterise Th1 (CD3+CD4+CD45RA-CXCR3+CCR6-), Th1* (CD3+CD4+CD45RA-CXCR3+CCR6+), and Th17 (CD3+CD4+CD45RA-CXCR3-CCR6+).**(d)** Gating strategy of follicular-like T helper cells (TfH) (CD3+CD4+CD45RA-CXCR5+). **(e)** Percentage of HLA-DR+ CD4+ T cells and **(f)** CD25 expression on CD4+ T cells in healthy controls and patients. **(g)** Gating strategy and assessment of % CD8+ T cells in *IVNS1ABP* patients (P) and healthy controls (C). **(h)** Naïve (CD19+IgD+CD27-), marginal-zone like (CD19+IgD+CD27+), and class-switched (CD19+IgD-CD27+) B cells in healthy controls (C) and *IVNS1ABP* patients (P). Two-sided Mann Whitney U. Bars represent mean ± S.E.M. Controls n=20, Patients n=4.

#### b) *ARPC1B* compound heterozygous patient

The index case presented aged 27-years-old with human papilloma virus associated cervical and vulval intraepithelial dysplasia grade 3. She had a past medical history of atopic dermatitis with recurrent staphylococcal skin infection, bacterial pulmonary and hepatic abscess that had necessitated intravenous antibiotics. She was the youngest of 10 siblings, and 2 older siblings had died during infancy. Immunological investigations revealed a raised IgE of 3200iu/ml and thrombocytopenia with a platelet count of $97x10^9/L$. During the follow-up period she developed anaplastic large cell lymphoma and was treated to complete remission with 6 cycles of cyclophosphamide, doxorubicin, vincristine and prednisolone (CHOP). Post-chemotherapy she suffered fatal influenza A and coronavirus pneumonitis causing respiratory failure.

Whole genome sequencing of the index case identified two heterozygous variants in *ARPC1B*, a short frameshift deletion leading to a premature termination codon (chr7:g.98988832_98988836del; ENST00000451682 c.739_743del; p.Leu247Glyfs*25) and a 9.1kb deletion spanning the promoter region of the gene (chr7:g.98967377-98976510del). Sanger sequencing of the variants in both parents and clinically unaffected siblings confirmed this to be a compound heterozygous variant in the patient (**Figure 3a**). The data confirming the variant contribution to ARPC1B deficiency from each variant is shown in **Figure 3b-d**. The single stain images of the patient and control monocyte derived macrophages vinculin is detectable in the patient’s cells; it does not encircle phalloidin dense structures, as it does in the controls, indicating a deficiency in podosome formation (**Figure 3e**).

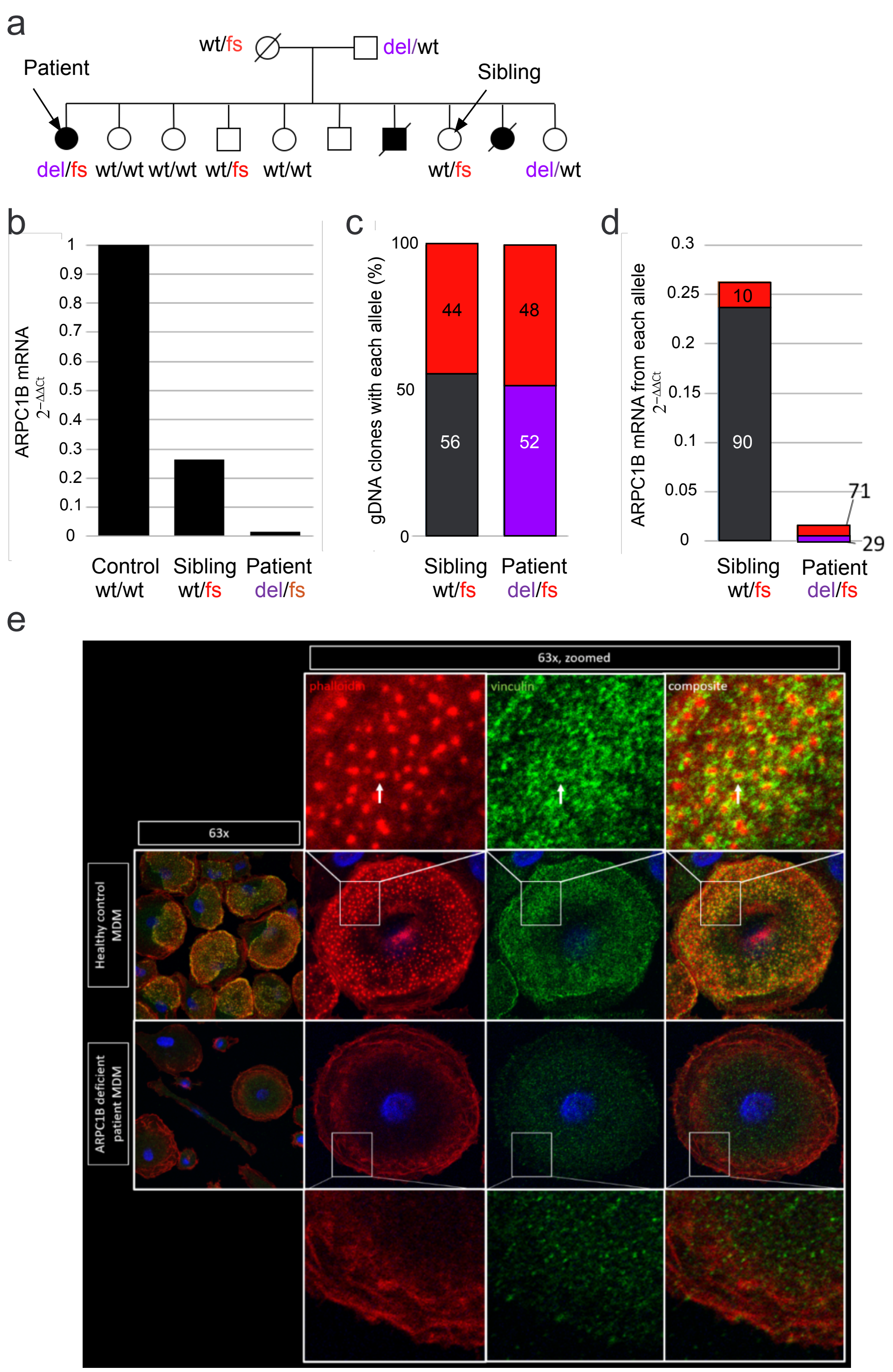


**Figure 3. Allele specific expression and Podosome formation in the ARPC1B patient. (a)** Pedigree of ARPC1B mutation patient and cosegregation of *ARC1B* genotype (wt- wild-type, del- deletion, fs – frameshift). **(b)** Histogram showing ARPC1B mRNA levels in healthy control, the patient and her sibling highlighted in (a). n=3 technical repeats **(c)** Allele-specific expression assay showing the ratio of wt, del and fs alleles in genomic DNA (gDNA) from peripheral blood mononuclear cells of the patient and sibling. n=3 technical repeats. **(d)** Relative expression of *ARPC1B* mRNA from each allele in the patient and sibling. Allele-specific expression assessed in complementary DNA (cDNA; synthesised from pre-mRNA). n=3 technical repeats **(e)** Podosomes were identified by staining adherent, fixed monocyte-derived macrophages for vinculin, phalloidin and the nuclear stain DAPI. Quantification was performed by counting podosomes on at least 100 cells per sample from 10 fields of view at 63x magnification from one independent experiment.

#### c) *SOCS1* and *PTPN2* cases

##### *SOCS1* patient

The index case is a 35-year-old female who presented with recurrent bacterial infections. She suffered repeated ear infections and aged 8-years-old requiring grommets. Aged 14 she suffered with bacterial pneumonia and investigation at this time showed panhypogammaglobulinaemia leading to a diagnosis of CVID and she was commenced on immunoglobulin replacement, then aged 9-years-old she suffered ITP requiring steroid treatment, high dose immunoglobulin, and anti-D treatment. Aged 22-years-old she developed chronic granulomatous uveitis which required continuous steroid eye drops. Her liver function tests were noted to be raised so a liver biopsy was performed showing an acute hepatitis. Aged 25 she developed a dry non-productive cough. CT chest showed areas of ground-glass with nodularity. Bronchoscopy did not identify any organisms and she underwent lung biopsy from which the histology showed florid lymphocytic infiltration with germinal centre formation and non-caseating granulomata consistent with granulomatous lymphocytic interstitial lung disease (GLILD). Treatment with daily 60mg prednisolone was commenced and lead to stabilisation, but no improvement in lung function. The steroid treatment concurrently improved her liver function tests suggesting an underlying autoimmune cause for hepatitis. At 29 years old she suffered a further severe relapse of ITP requiring treatment rituximab improved her uveitis, allowing cessation of steroid eye drops, and resolved the much of the GLILD seem on HRCT chest imaging.

Whole genome sequencing identified the novel frameshift variant in *SOCS1* (chr16:g.11348855insGCGGC; ENST00000332029 c.480_481insGCGGC; p.Met161Alafs*46), and Sanger sequencing of parents' DNA confirmed it to be de-novo (**Figure 4a**). None of the relatives have a history suggestive of immunodeficiency or autoimmunity.

Additional unrelated cases of *SOCS1* variants were identified with GeneMatcher[3]. B.II.1 is a 14-year-old female who presented at the age of 9-years-old with recurrent *streptococcus pneumoniae* pneumonia which led to the development of a pneumatocele, and repeated cutaneous herpes zoster outbreaks. At the age of 10-years-old she developed eczema and alopecia areata. This was followed by further autoimmune phenomena between the ages of 11-years-old with Evan's syndrome (ITP, autoimmune haemolytic, and autoimmune neutropenia). She remains on antimicrobial treatments as needed and thrombopoietin for chronic ITP. Whole exome sequencing identified the novel heterozygous stop-gain variant *SOCS1* ENST00000332029 c.192C>G; p.Tyr64*. Her father B.I.1 was also found to have the same p.Tyr64* variant (**Figure 4b**). He has a long history of autoimmune Hashimoto's thyroiditis and recurrent jaundice due to liver disease. Recently he has developed non-infective colitis with chronic diarrhoea which is currently under investigation. Western blot demonstrated reduced SOCS1 and prolonged phosphorylation of STAT1 in patient T cell blasts compared with health controls **(Figure 4c)**.

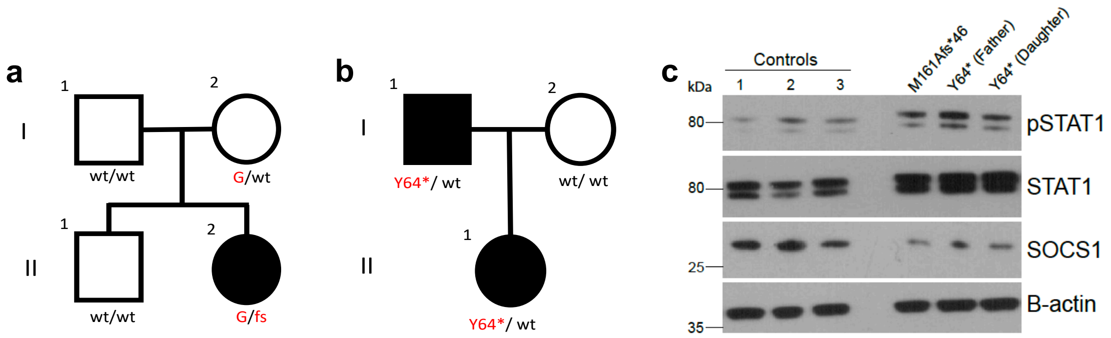


**Figure 4. Pedigrees of the *SOCS1* patients. (a)** The pedigree diagram shows the co-segregation of the rare de-novo frameshift variant p.Met161Alafs*46 and the common risk polymorphism rs2286974-G (G) within the family. **(b)** Pedigree of Family B showing segregation of the novel stop-gain p.Tyr64*. **(c)** Western blot of controls n=4 and affected SOCS1 patients of SOCS1, STAT1 and phosphorylated STAT1 (pSTAT1) 2 hours after stimulation with IFNγ. Blot representative of 3 technical replicates. For gel source data, see Supplementary Figure 1.

#### *PTPN2* family

The male index case presented age 20-years-old for evaluation with a 2-year history of recurrent bacterial sino-pulmonary infections and resultant bibasal bronchiectasis. Immunological investigations showed panhypogammaglobulinaemia and low B cells, and he was diagnosed with common variable immunodeficiency (CVID). He was commenced on immunoglobulin replacement and had a marked reduction in bacterial infectious burden. His past medical history consisted of severe deforming arthritis in a rheumatoid distribution that started aged 9, and which had resulted in bilateral ulnar deviation. The arthritis was treated with long courses of prednisolone monotherapy and entered remission at the age of 18 years old and has remained quiescent in the absence of further therapy.

During the follow-up period of over 20 years he has suffered with granulomatous inflammation with massive splenomegaly and granulomatous lymphocytic interstitial lung (GLILD) disease. His GLILD is monitored by serial pulmonary function tests and he remains asymptomatic despite radiological and pulmonary function changes.

The mother of the index case is a 75-year-old female with a history of lupus diagnosed when she was 45 years old. This presented as pleurisy for which she required prednisolone treatment. Aged 42 she had been diagnosed insulin dependent diabetes mellitus. She also has a background history of neutropenia, hypothyroidism, and asthma. The patient's father passed away at the age of 73 from a brain tumour and has no known history suggestive of immunodeficiency or autoimmunity. Of the two siblings, his 46-year-old sister is healthy the brother has a history of recurrent mucosal polyposis and chronic sinusitis.

Whole genome sequencing of the index case revealed a novel heterozygous stop-gain variant in *PTPN2* (chr18:g.12802138C>A; ENST00000309660 c.871G>T; p.Glu291*) which was subsequently found to be present in his mother (**Figure 5a**). The index case also carries a common *PTPN2* variant rs2847297 (chr18:g.12797694A>G), subsequently found to be shared with his brother but not the mother; therefore, the common variant must have been inherited from the father, and the index case is compound heterozygous for the two variants (**Figure 5a**). The common variant rs2847297 has been shown to be associated with several autoimmune conditions. Allele specific expression assay demonstrated a marked reduction in expression from the rs2847297 (**Figure 5b**).

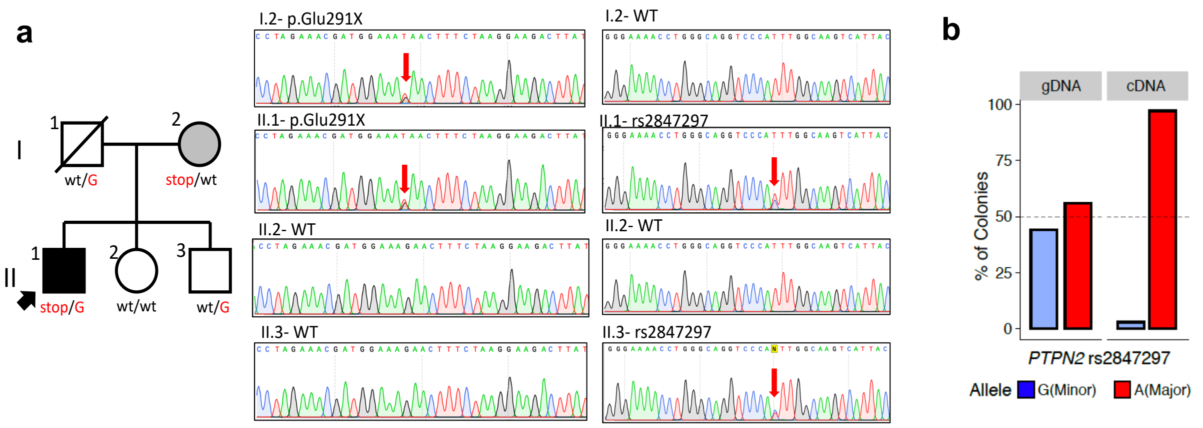


**Figure 5. Pedigree of the *PTPN2* kindred. (a)** The pedigree diagram demonstrates co-segregation of the intronic expression quantitative trait loci (eQTL) rs2847297-G with the stop variant Glu291*. Sanger sequence traces confirm the genotypes in each family member. **(b)** Allele specific assay from DNA of II.3 (patient's brother) showing amounts of gDNA and reverse transcribed cDNA. n=2 technical repeats

#### *PTPN2 and SOCS1* Immunophenotype

Whilst mice deficient in *Ptpn2 an Socs1* both die from fatal Type 1 inflammation associated disease, this occurs slightly earlier in the *Socs1* knock-out at 2-3 weeks[4] than in the *Ptpn2* knock-out[5] at 3-5 weeks. Whilst an inflammatory phenotype has been described in the haploinsufficient *Socs1* mouse[6,7] this has not been described *for Ptpn2* haploinsufficient mice.

Although both the *PTPN2* Index case and their mother have the *PTPN2* protein truncating variant, only the *PTPN2* index case has the *PTPN2* haplotype (on their other allele) associated with reduced *PTPN2* transcription (**manuscript Fig. 4d**). The *SOCS1* and *PTPN2* patients show low/normal CD4+ T cells and low/normal B cells (**Figure 6a-b**). Additionally, the *SOCS1* and *PTPN2* cases show a Th1 skew and lower Treg numbers (**Figure 6d-e**) as observed in the *SOCS1* haploinsufficient and homozygous knock-out *PTPN2* mice[6,8].

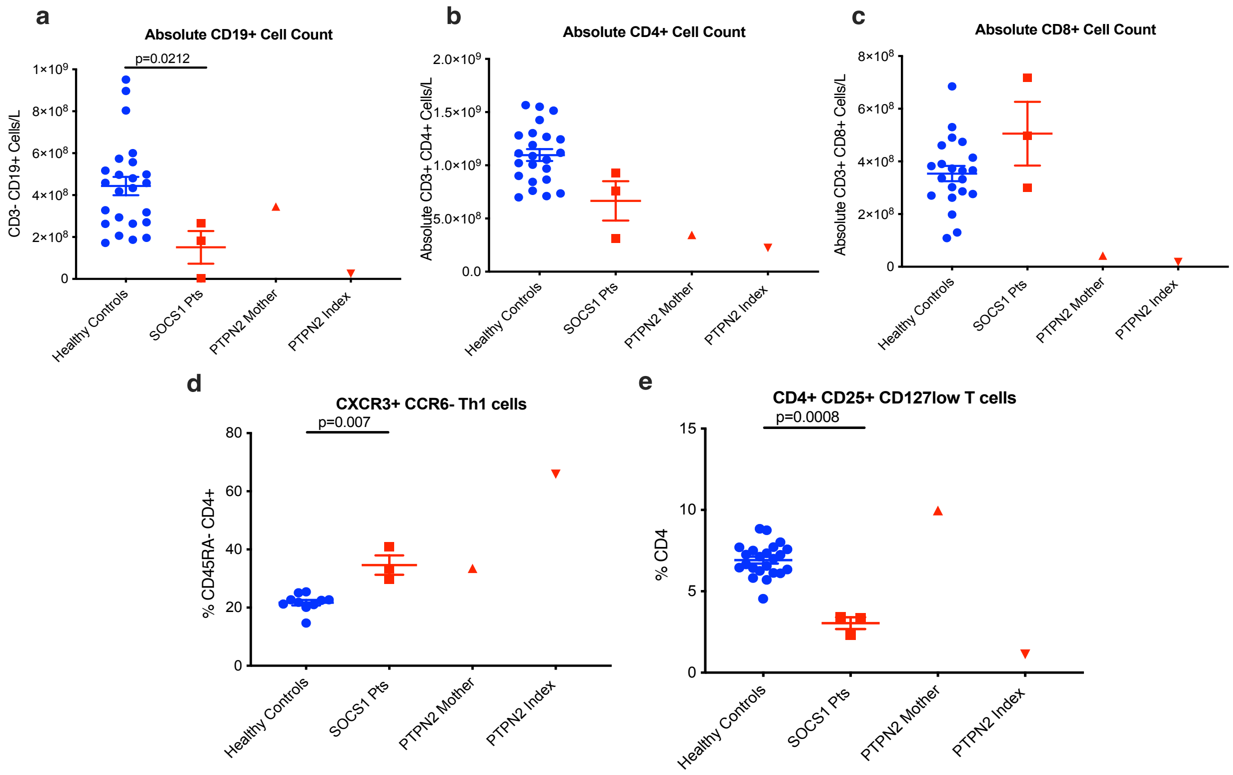


**Figure 6. B and T cell phenotypes of SOCS1 and PTPN2 families. (a)** Absolute B cell (CD19+) count in cells/L in healthy controls (n=24) (blue) compared with SOCS1 patients (n=3), and the PTPN2 mother and index cases. **(b)** Comparisons of absolute CD4 (CD3+CD4+) T cell count in cells/L (controls n=23), and **(c)** absolute CD8 (CD3+CD8+) T cell count in cells/L (controls n=21). **(d)** Proportion of CD45RA- CD4+ T cells which are Th1 cells (CXCR3+ CCR6-) in healthy controls (n=10) and SOCS1 patients (n=3), and the PTPN2 mother and index cases. **(e)** Proportion of CD4+ T cells which are regulatory T cells (CD25+ CD127low) in healthy controls (n=23) and SOCS1 patients (n=3) and the PTPN2 mother and index. All tests two-sided Mann-Whitney U test. Bars mean ± S.E.M.

### Supplementary Note 2

### Population stratification in the PID cohort

Cryptic population stratification is recognised as a potential source of false positive associations when exploring rare variants in the context of GWAS[1]. It is therefore necessary to define and minimize its potential impact on causal gene prioritization using BeviMed.

#### 1. Removal of relatives

The detailed method for identifying related individuals and deriving the maximal set of unrelated individuals among the 13,037 participants in the NBR-RD cohort is described in[2]. Briefly, we first generated a set of 32,875 high-quality genome-wide SNPs and their genotype matrix was encoded in the merged VCF file. We then used PC-AiR[3] to obtain principal components (PCs) based on these genotypes, and the PC-Relate[4] function to compute a kinship matrix that accounts for the population structure captured by the leading 20 PCs. We passed this kinship to the PRIMUS[5] function to obtain clusters of related individuals. We defined a maximal set of unrelated individuals based on a pairwise kinship coefficient <=0.09.

We then identified a set of 2,110 unrelated non-admixed individuals in the 1000 Genomes Phase 3 data, merged their genotypes with those of the 13,037 individuals in our cohort, and ran the principal component analysis (PCA). Because the 1000 Genomes collection is genetically diverse, combining it with our sequence dataset meant that a greater amount of genetic diversity was captured by the PCA. The whole pipeline generated a maximal set of 10,259 unrelated individuals which was then used for case-control analyses for cohort-wide allele frequency estimation.

#### 2. Population structure

As allele frequency datasets for non-Europeans are much smaller than for Europeans, potential false positives may be induced by the unintentional inclusion of variants that appear rare but occur in greater frequency in populations of non-European ancestry. Unrelated individuals were partitioned into non-Finnish Europeans, Finns, Africans, South Asians, East Asians and 'Other' using 1000 genomes population code annotations, as described in [2]. Briefly, we projected sample genotypes onto a PCA basis assembled from the leading five components from 1000 genomes and used multivariate normal distributions to compute the likelihood that a given sample belonged to a given ancestral category. Samples that could not be confidently assigned to any of these categories were labelled 'Other'.

To guard against associations driven solely by ethnicity, we implemented the following quality control process: 1 – run BeviMed to obtain individual gene PPA; 2 – if PPA >0.1 (signifying potential gene of interest with >10% posterior probability of being causal), re-run BeviMed after excluding all variants present in heterozygosity in at least two individuals with the same non-European ancestry (e.g. South Asian or African) and absent from all Europeans (homozygosity in a non-European case is not in itself considered to be indicative of a high allele frequency because it is much more likely the result of consanguinity); 3 – if the new BeviMed PPA <0.1, flag the result as being dependent on variants from a non-European population. Out of the top 50 prioritised genes only two, *TMEM129* and *ZC3HC1*, dipped below 0.1 and were therefore flagged by this procedure (**Figure 1** and **Figure 2** below**;** see also **Supplementary Table 2, manuscript Fig. 2a**).

#### 3. Rare variant cryptic relatedness

Whilst BeviMed association analyses exclude closely related individuals, there exists the possibility that some might be distantly related and share rare variants, which could give rise to the prioritisation of spurious genes. Of the top 50 BeviMed prioritised genes (**Supplementary Table 2**), 29 were driven by variants that were not shared between individual cases and therefore cannot be the result of cryptic relatedness.

It is not unexpected that, for the remaining 21 prioritised genes, some cases shared rare variants. Such sharing of variants is common in PID (see **manuscript Fig. 1c**), and is thought to be driven at least in part by the selection of such variants because they specifically drive pathology. To address the possibility that cryptic relatedness might have contributed to the prioritization of some of these, we first examined whether there was additional sharing of rare variants for such cases outside of that observed within the BeviMed prioritised gene; a strong indication of cryptic relatedness. This analysis identified such sharing in 9 of the 21 sets of cases (*STAT1*, *NFKB2*, *ZFP36, ODC1*, *G6PC3, RN7SL142P*, *TMEM132D, METTL2B* and *CTD-2010I22.2)*.

For the residual pairs of cases where only a single rare variant within the prioritised gene was shared, we sought further evidence of cryptic relatedness. If the sharing of the variant between a pair of individuals is being driven by cryptic relatedness, we would expect that they will also share a common genetic background at that locus; if on the other hand the variant has arisen independently, such sharing will be absent. To investigate this we used TRUFFLE[6] (--segments --ibs1markers 4000 --ibs2marker 500) to identify, using unphased common variant genotypes (MAF>5%), chromosomal segments that were identical by descent across affected individuals harbouring a single shared ultra-rare variant for the remaining 15 BeviMed prioritised genes. We found identical by descent chromosome segments overlapping the shared ultra-rare variant in two additional genes (*ZNF34* and *SLC13A4*). Of the 13 genes flagged by these procedures (**Figure 2** below; see also **Supplementary Table 2**, **manuscript Fig. 2a**), at least 3 are IUIS `known' genes (*STAT1*, *NFKB2* and *G6PC3*) and thus whilst such cryptic relatedness could lead to erroneous prioritisation, it does not automatically preclude PID causal gene candidacy.

Additionally, we performed two SKAT analyses to model population stratification: one with and one without the top 10 PCs as covariates. Correcting for 10 PCs did not change the enrichment of known PID genes amongst the most highly ranked 76 (the number of BeviMed results with a posterior probability of association greater than 0.1) results from the SKAT analysis (Fisher's exact test p-values: SKAT without adjustment: $4.7x10^{-6}$; SKAT with adjustment for 10 PCs: $4.7x10^{-6}$; BeviMed: $3.1x10^{-8}$). Amongst the top 50 BeviMed ranked genes, there were only three for which signal strength was significantly decreased once the 10 PCs were included in the model and could be the only additional potential "false positives": *ARPC1B* (a known PID gene), *ZC3HC1* (already flagged by cryptic relatedness procedure), and *ZFAND3* (BeviMed rank 38).

In summary, an effect of population stratification on some genes prioritised by BeviMed cannot be excluded. We have, however, taken measures to minimise and measure this, and can be confident that it is not having a substantial impact on gene prioritization. To make this clear we have clearly labelled the 14 genes flagged by these analyses (2 through non-European ancestry and 12 though cryptic relatedness) in **Supplementary Table 2** and **Figure 1** below. Provided any effect of population stratification in driving rare variant associations is relatively minor, it does not harm the Bayesian approach to causative gene discovery, but it does underline the important point that genes identified by BeviMed cannot be considered causative without further functional validation.

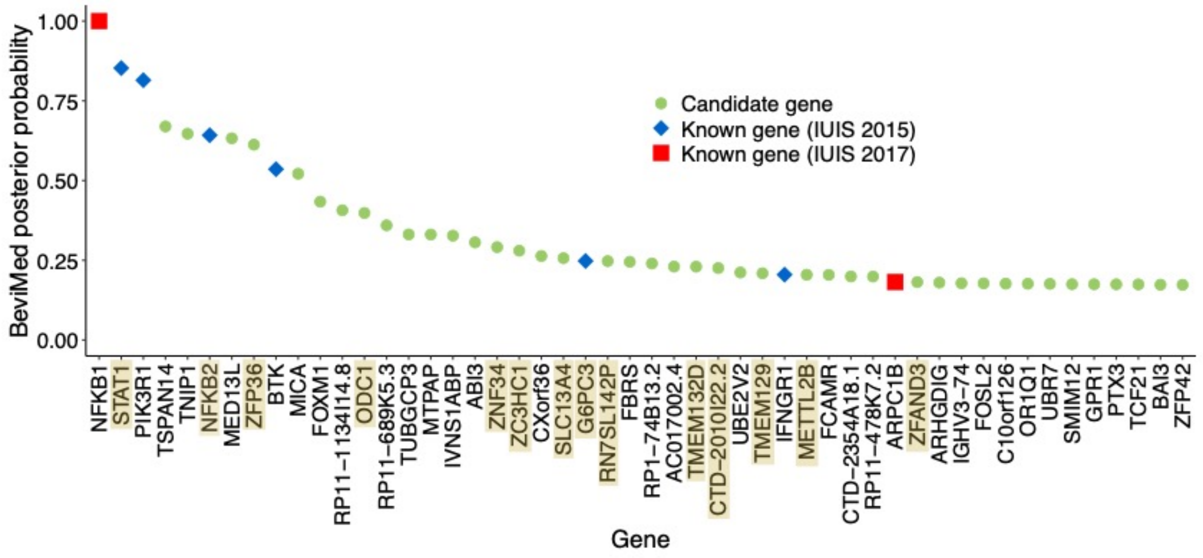


**Figure 1. The top 50 candidate PID genes prioritised by BeviMed analysis.** Highlighted in yellow are those identified in the population stratification analyses described here as potentially confounded by population stratification. The analysis was performed on 886 PID index cases and 9,284 unrelated controls.

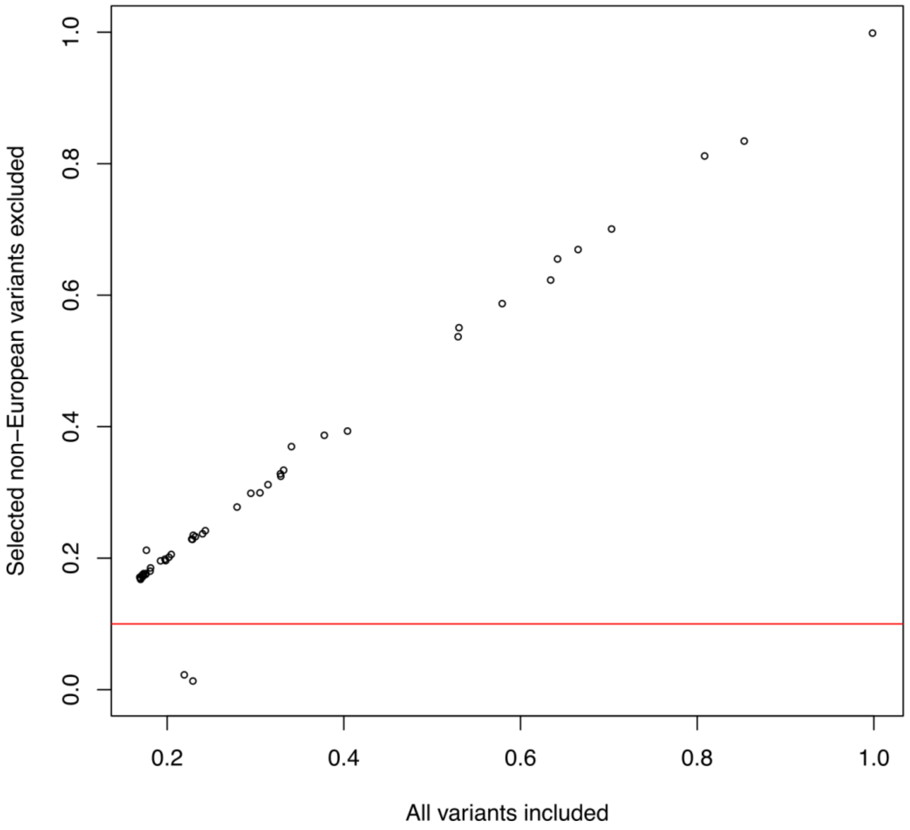


**Figure 2. Flagging of genes with high BeviMed posterior probabilities of association (PPA) due to inclusion of variants in non-European populations.** Individual data points represent PPA for genes shown in Figure 1 above and Supplementary Table 2 (cases n = 886; controls n = 9,284). Initial results based on all rare variants selected for analysis with BeviMed (x-axis) are plotted against new results obtained after excluding variants according to the quality control process for flagging genes with initially high PPA due variants that are only observed in a non-European population (y-axis), and whose new PPA dips below the specified threshold of <0.1 (red line). The two genes with new PPA<0.1 and flagged by this procedure are *ZC3H1* and *TMEM129*.

### Supplementary Note 3

### AD-PID GWAS analysis

#### Mitigating sources of inflation in association statistics

Given the `real world' nature of the NBR-RD cohort, it was not possible to select ancestry-matched controls when performing the AD-PID GWAS, which might result in spurious associations due to population stratification. We, therefore, employed a number of strategies in order to mitigate population stratification and other technical effects on our GWAS analysis:

<u>Technical confounders:</u> Cases and controls were sequenced across a 4-year period using 3 different Illumina chemistries and read lengths. These were controlled for by including read length as a covariate during association testing.

<u>Population stratification:</u>

- We conducted PCA analysis on the genotype matrix (**Figure 1**), and used the first 10 principal components as logistic regression covariates in order to minimise the effects of population stratification on downstream analysis.
- A quantile-quantile normal plot (**Figure 2**) of GWAS P-values showed residual inflation of association statistics indicating the presence of a small amount of residual population stratification (lambda=1.022). To account for this we used the Genomic Control method in order to adjust P-values.
- To account for population stratification that might be present in *Li et al.*, we again applied the Genomic Control method using a lambda of 1.039[1], prior to performing any meta-analysis.

Alternatively, to confirm the robustness of the approach described above we performed an additional AD-PID GWAS analysis, with the same covariates, using the BOLT-LMM software[2], which employs a linear mixed model approach in order to control for confounding due to population stratification and cryptic relatedness at the individual SNP level. After genomic control we found that differences between the two approaches were minimal (**Table 1**) and therefore took forward our initial results for further analysis.

| | Uncorrected | | | Genomic Control | | | BOLT-LMM+Genomic Control | |
|---|---|---|---|---|---|---|---|---|
| SNP | AD-PID P | Li *et al.* P | Meta P | AD-PID P | Li *et al.* P | Meta P | AD-PID P | Meta P |
| Genome-wide significant signals (MAF > 5%) | | | | | | | | |
| rs2517529 | 8.60E-10 | 8.80E-11 | 4.30E-19 | 1.30E-09 | 1.96E-10 | 1.50E-18 | 9.18E-10 | 1.02E-18 |
| rs2286974 | 4.90E-06 | 3.40E-08 | 9.60E-13 | 6.23E-06 | 6.18E-08 | 2.20E-12 | 4.08E-06 | 1.50E-12 |
| Additional suggestive signals (MAF > 5%) | | | | | | | | |
| rs3806624 | 2.20E-03 | 2.20E-06 | 3.20E-08 | 2.43E-03 | 3.41E-06 | 5.30E-08 | 2.46E-03 | 6.17E-08 |
| rs12563449 | 7.30E-02 | 1.10E-08 | 6.90E-07 | 7.66E-02 | 2.13E-08 | 1.37E-07 | 5.80E-02 | 1.29E-06 |
| rs80191532 | 6.90E-07 | 4.60E-02 | 6.90E-07 | 9.14E-07 | 5.06E-02 | 9.30E-07 | 1.38E-06 | 1.98E-06 |
| rs10750403 | 3.70E-03 | 4.00E-04 | 4.90E-06 | 4.07E-03 | 5.13E-04 | 7.00E-06 | 3.13E-03 | 5.31E-06 |
| rs11851820 | 1.20E-02 | 1.30E-08 | 4.90E-06 | 1.29E-02 | 2.47E-08 | 7.10E-06 | 1.30E-02 | 2.60E-05 |
| Additional AD-PID genome-wide significant signal (MAF>0.5%) | | | | | | | | |
| rs34557412 | 7.87E-13 | N/A | N/A | 1.37E-12 | N/A | N/A | N/A | N/A |

**Table 1. Comparison of methods for adjusting for population stratification across lead index variants listed in Extended Data Table 3.** Uncorrected and Genomic Control corrected test statistics are based on single variant logistic regression, using the first 10 ethnicity analysis principal components as covariates. BOLT-LMM linear mixed-model association test statistic is based on residuals from Bayesian modelling. AD-PID GWAS details: cases n=733, controls n=9,225, lambda=1.022. Li *et al.* GWAS details: cases n=778, controls=10,999, lambda=1.039.

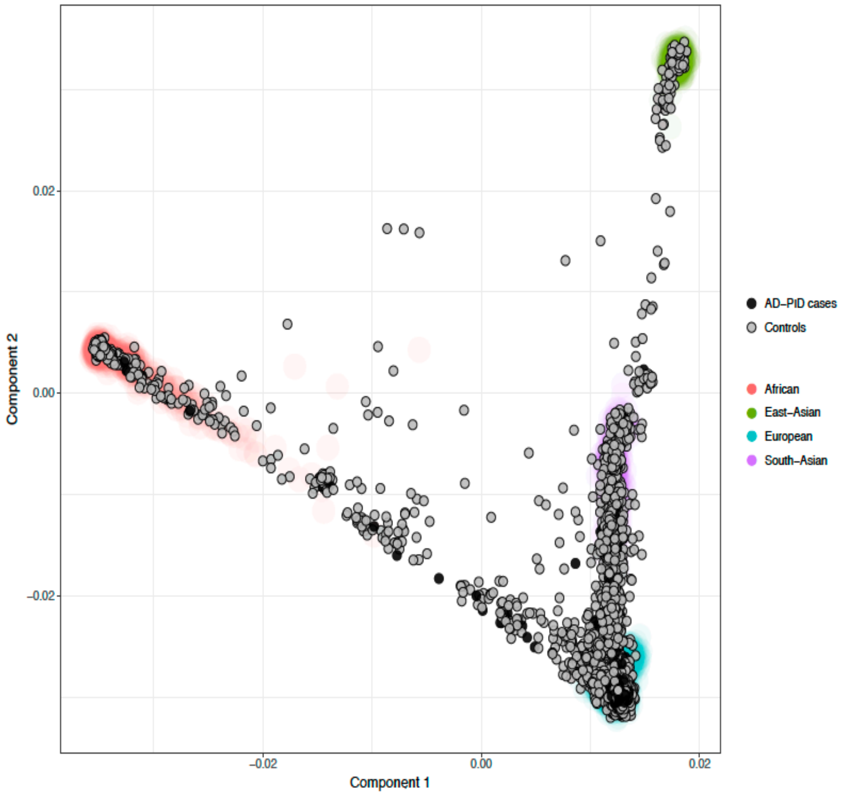


**Figure 1. Principal component analysis of AD-PID case (n=733) and control (n=9,225) samples.** The plot shows the projection of the genotype matrix of cases (black circles) and controls (grey circles) onto the 1000 Genomes derived PCAs. The 1000 Genomes samples are shown as diffuse points underneath in colour.

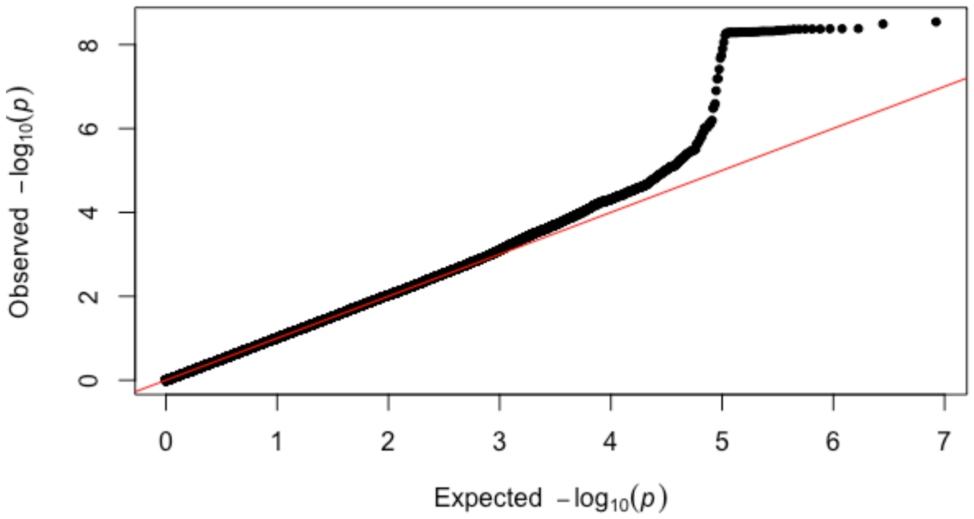


**Figure 2. A Quantile-Quantile normal plot of AD-PID association P values (Lambda = 1.022).** The *x* and *y* axes show the expected and observed logistic regression -log10(P) for the AD-PID GWAS respectively (AD-PID cases = 733; controls = 9,225). The red line shows *x*=*y*.

##### 6p21.33: MHC locus

Conditional analysis of lead SNPs in the AD-PID GWAS suggested two independent signals (**Extended Data Fig. 7a**) that localise to the HLA class I and class II regions. We therefore imputed the HLA classical alleles and amino acid polymorphisms in HLA genes and repeated the association and conditional analyses.

Imputation of HLA classical alleles showed the strongest associations with class I HLA-B*08:01 and class II HLA-DQA*01:02 alleles, but neither signal reached genome-wide significance on its own or when conditioning on the other allele (**Extended Data Fig. 7b**). Analysis of the imputed amino acids at multi-allelic positions of the MHC locus genes identified the strongest associations with Asparagine at position 114 of *HLA-B* (AA_B_114_31432129_N) and Arginine/Lysine at position 71 of *HLA-DRB1* genes (AA_DRB1_71_32659935_RK) (**Extended Data Fig. 7c**). Both *HLA-B* Arg114 and *HLA-DRB1* Lys71 signals were genome-wide significant and remained so upon conditioning on each other, consistent with independent effects. Conditioning on both residues removed all of the association signal at this locus (bottom panel in **Extended Data Fig. 7c**).

In the context of HLA-B, changes at position 114 have been shown to alter peptide binding[3]. The His114Asn polymorphism alters the depth of the antigen binding groove of class I allele HLA-B*0801, thus promoting the presentation of peptides of various sizes that are necessary for effective T cell responses to invading pathogens[4]. Asn114 polymorphism has been found to be more associated with HIV control than the ancestral HLA-B*08:01 allele with Histidine residue at this position, and this association was shown to be driven by altered peptide presentation[5]. Likewise, amino acid position 71 in HLA-DRB1 has been shown to be critical for peptide presentation by HLA-DR molecules[6].

#### 3p24.1: *EOMES*

An AD-PID association at 3p24.1 (**Figure 3**) indexed by rs3806624 was identified. The same SNP has been associated with the autoimmune disease, rheumatoid arthritis, in a previous meta-analysis[7]. Although initially defined promoting CD8+ T cell effector function[8], other studies have highlighted its role in preventing terminal differentiation[9]. In CD4+ T cells, downregulation of *EOMES* by *IRF4* is required for the commitment to a T follicular helper (Tfh) cell phenotype, as opposed to a T helper (Th)1 cell differentiation following interleukin (IL)12 stimulation of naïve T cells (Th0)[10]. In humans, common *EOMES* variants have been associated with lymphocyte counts[11], Hodgkin's lymphoma[12], multiple sclerosis[13], and rheumatoid arthritis[7] from GWAS data (**Table 2**). The association of this variant rs3806624 in PID and also with autoimmunity provides further evidence of the case biological similarities between PID and autoimmunity.

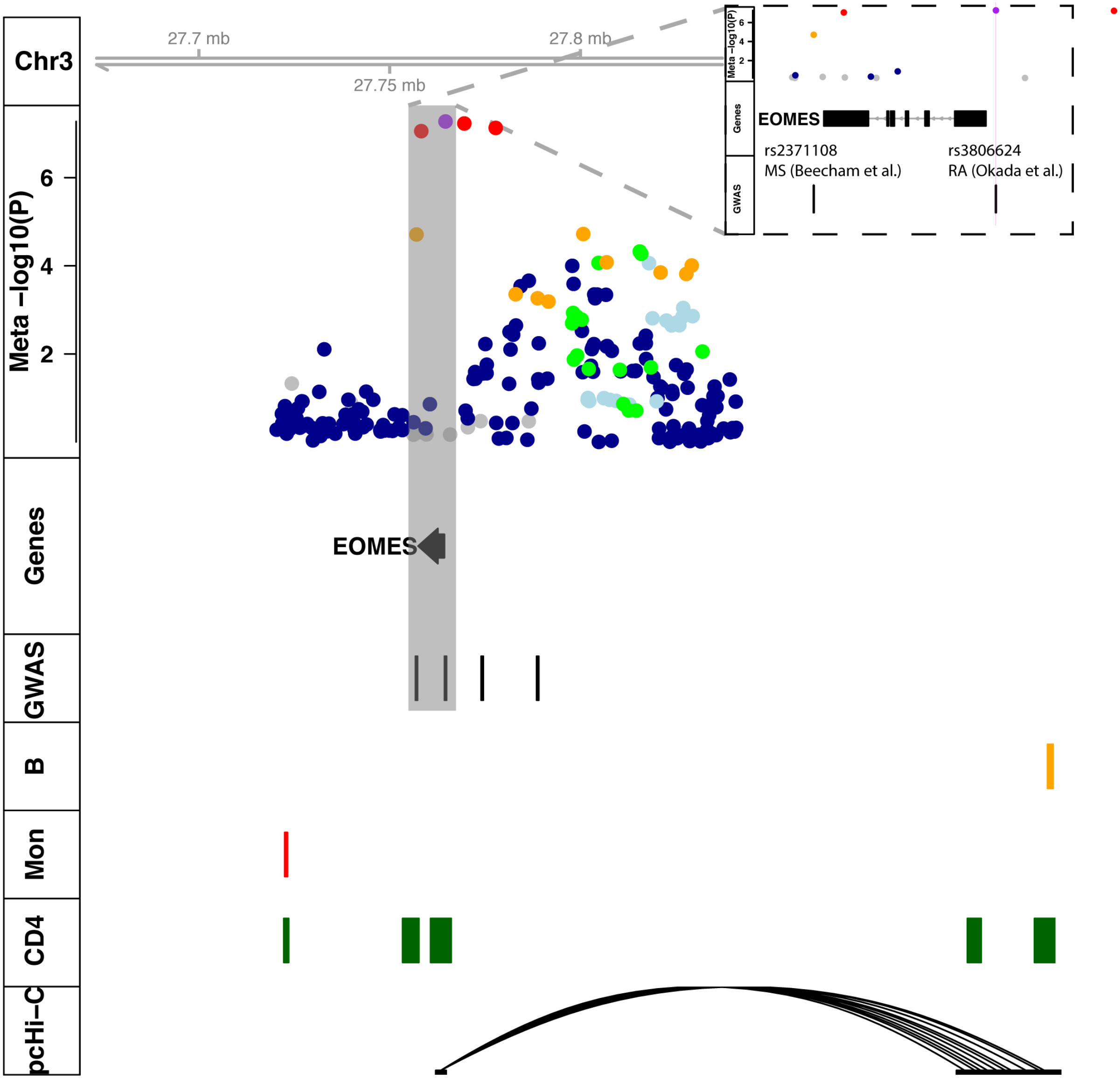


**Figure 3. Regional AD-PID meta-analysis association plot of 3p24.1 (*EOMES*) region.** Tracks are as follows: **Meta -log(P)** dot plot of AD-PID fixed effects association meta-analysis with Li *et al.* (total cases n=1,511; controls n=20,224). The index SNP is purple, others are coloured based on LD information from UK10K project with red indicating high LD (r^2>0.9), blue low LD (r^2<0.2) and grey where no information available. **Gene** - Canonical gene annotation (Ensembl V75), **GWAS** - location of index variants from other immune-mediated diseases, **CD4**, **B**, **Mon** - putative regulatory regions in CD+T-cells, Total B cells and Monocytes computed from the union of ATAC-Seq and H3K27ac ChIP-Seq data, **pcHi-C** - Promoter Capture Hi-C interactions, in above primary cell types, with the exception of **Meta -log(P)** which shows dot plot of AD-PID association meta-analysis with Li *et al.* Detail shows the location of RA index SNP that overlaps AD-PID index variant and its promoter proximity.

| SNP | Disease/ Trait | Study sample size | P-value | $r^2$ with rs3806624 |
|---|---|---|---|---|
| rs11129295 | Multiple sclerosis[13] | case n=9,772; ctrl n=17,376 | 1.00E-09 | 0.67 |
| rs2371108 | Multiple sclerosis[14] | case n=14,498; ctrl n=24,091 | 2.00E-15 | 0.72 |
| rs3806624 | Rheumatoid arthritis[7] | case n=29,880; ctrl n=73,750 | 3.00E-08 | 1.00 |
| rs3806624 | Hodgkin's lymphoma[12] | case n=1,465; ctrl n=6,417 | 1.00E-12 | 1.00 |
| rs6801231 | Lymphocyte count[11] | total n = 171,643 | 7.00E-13 | 0.67 |

**Table 2. GWAS Catalogue (https://www.ebi.ac.uk/gwas/) index variants for other published genome-wide significant associated traits at the 3p24.1 locus.** See referenced studies for full details of association tests used to generate the quoted P-values. The index variant for our meta-analysis is rs3806624.

##### 16p13.13: *SOCS1*

An AD-PID association at 16p13.13 (**Figure 4**), indexed by rs2286974, within intron 19 of *CLEC16A* was identified. This locus has been previously found to be associated with multiple immune-mediated diseases including CVID[1], Type 1 diabetes[15], Celiac disease[16], Primary Biliary Cirrhosis[17], Psoriasis[18], Multiple sclerosis[14], SLE[19] and IBD[20] although there is limited evidence that causal variants are shared[19]. Functional work has suggested putative functional roles for *CLEC16A* in CVID, *DEXI*[21] in T1D and MS. Integration of our AD-PID meta-analysis results with promoter capture Hi-C data for relevant primary human tissues[22] additionally implicated *PRM2*, *RMI2*, *PRM3*, *TNP2* and *SOCS1*. The biology of SOCS1 is discussed in the main text.

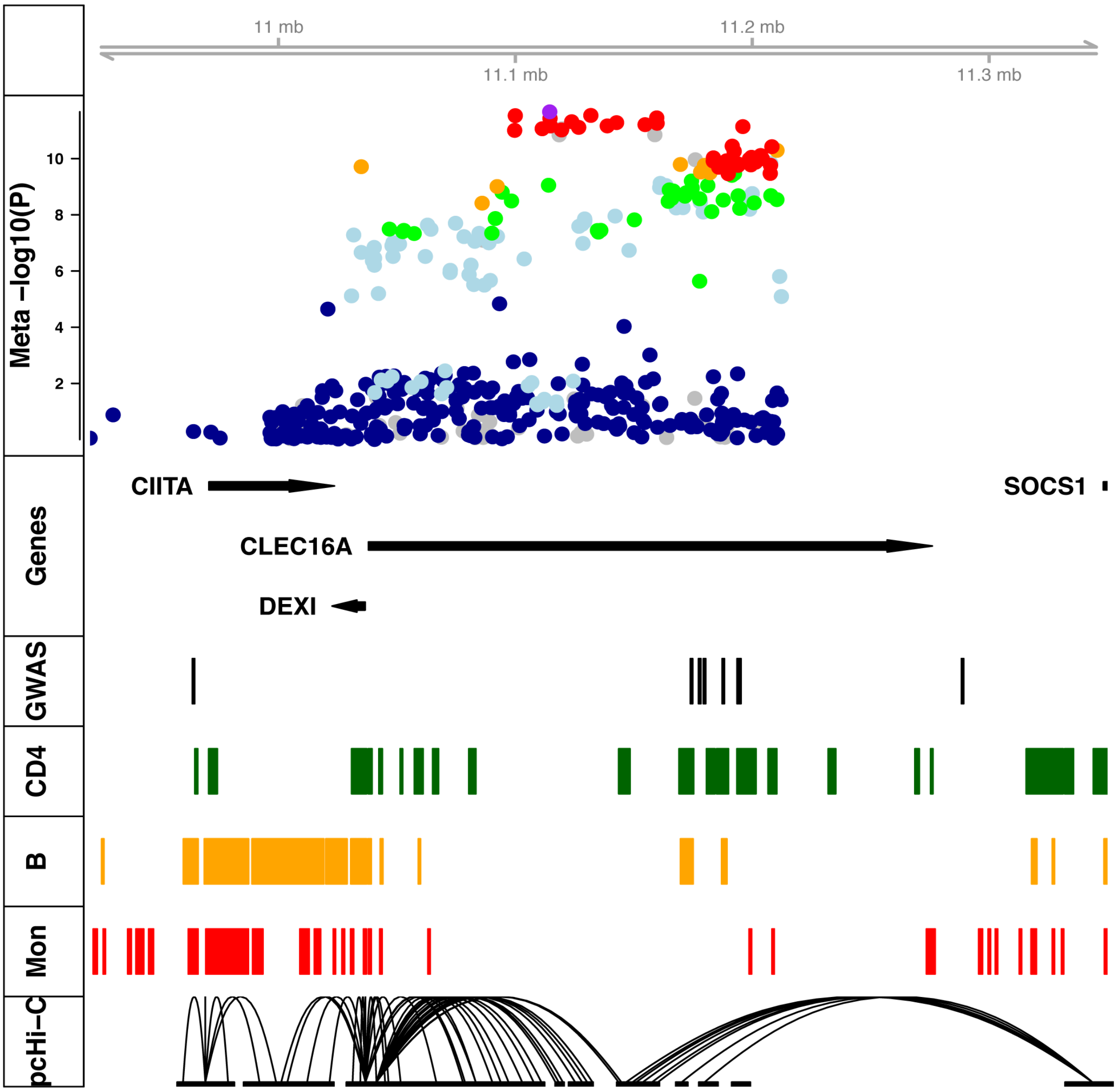


**Figure 4. Regional AD-PID meta-analysis association plot of 16p13.13 (*CLEC16A*/*SOCS1*).** Tracks as described in Figure 3. Tracks are as follows: **Meta -log(P)** dot plot of AD-PID fixed effects association meta-analysis with Li *et al.* (total cases n=1,511; controls n=20,224). The index SNP is purple, others are coloured based on LD information from UK10K project with red indicating high LD (r^2>0.9), blue low LD (r^2<0.2) and grey where no information available. **Gene** - Canonical gene annotation (Ensembl V75), **GWAS** - location of index variants from other immune-mediated disease, **CD4**, **B**, **Mon** - putative regulatory regions in CD+T-cells, Total B cells and Monocytes computed from the union of ATAC-Seq and H3K27ac ChIP-Seq data, **pcHi-C** - Promoter Capture Hi-C interactions, in above primary cell types, with the exception of **Meta -log(P)** which shows dot plot of AD-PID association meta-analysis with Li *et al.*

#### 18p11.21: *PTPN2*

Our data identified a single variant downstream of *PTPN2* as associated with PID (**Figure 5**). *PTPN2* encoded the T-cell protein tyrosine phosphate (TC-PTP) protein, a non-receptor tyrosine-specific phosphatase that dephosphorylates both receptor and non-receptor associated tyrosine kinases[23]. Genome-wide association studies have implicated *PTPN2* variants as associated with multiple autoimmune diseases including inflammatory bowel disease[24], rheumatoid arthritis[7], juvenile inflammatory arthritis[25], and type 1 diabetes[15]. *PTPN2* also has an important role in cancer. Somatic *PTPN2* deletions are present in up to 6% of T-cell acute lymphoblastic (T-ALL) leukaemia cases contributing to JAK1 oncogenicity, and aberrant expression of *TLX1* in T-ALL[26,27]. Somatic variants in *PTPN2* have also been identified in other malignancies with the germline *PTPN2* variant p.E291X, presented in this report is observed in cancer genome atlases[28]. TC-PTP negatively regulates many cell cytokine receptors and in the somatic context of colon cancer, *PTPN2* haploinsufficiency may contribute to enhanced epithelial growth factor receptor (EGFR) signalling and epithelial cell-derived cancer growth[29]. The immunobiology of *PTPN2* is discussed in the main text.

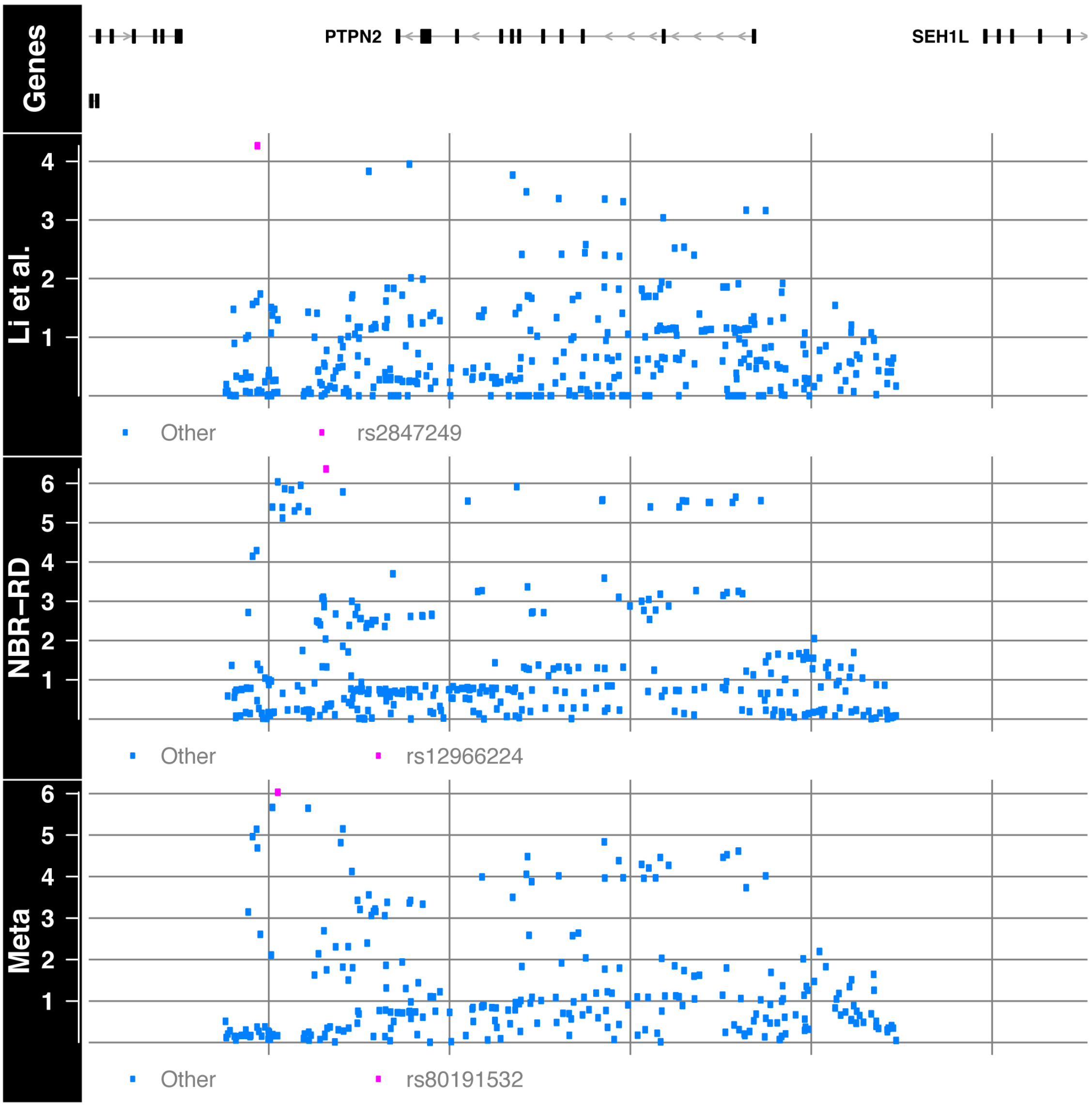


**Figure 5. Comparison of GWAS association signals at 18p11.21 for Li *et al.*, NBR-RD AD-PID (this study; cases n=733; controls n=9,225), and Meta (fixed effects meta-analysis; cases n=1,511; controls n=20,224).** Y-axis is -log10(P) of the univariate association statistic. Top SNP in each study is marked in cyan.

##### 17p11.2 *TNFRSF13B (TACI)*

A partially penetrant monogenic cause of CVID was described in 2005[30,31] after enrichment of variants within CVID cohorts were found within the coding region of *TNFRSF13B* (TACI) on chromosome 17. However subsequent follow-up studies clarified that these same variants are also found within the healthy population[32]. This study is the first to describe using a GWAS methodology, a genome-wide association signal implicating the previously reported rs34557412 coding variant and two non-coding variants in high LD with AD-PID susceptibility (**Figure 6**). The two non-coding variants align with cis-regulatory elements in relevant cell types, one of which is proximal to the promoter of *MPRIP*, furthermore, promoter-capture Hi-C evidence[22] links both variants through a possible chromatin interaction. This raises the possibility that the discrepancy between the murine knock-out and the human TACI variant phenotypes[30] may be attributable to the lead coding SNP not being solely responsible for modulating risk at this locus.

TACI, BAFFR and BCMA make up the three receptors for the ligands BAFF and APRIL. The interactions between these receptors and ligands is necessary for normal B lymphocyte differentiation into antibody-producing cells; positioning them as prime candidate loci for antibody deficiency

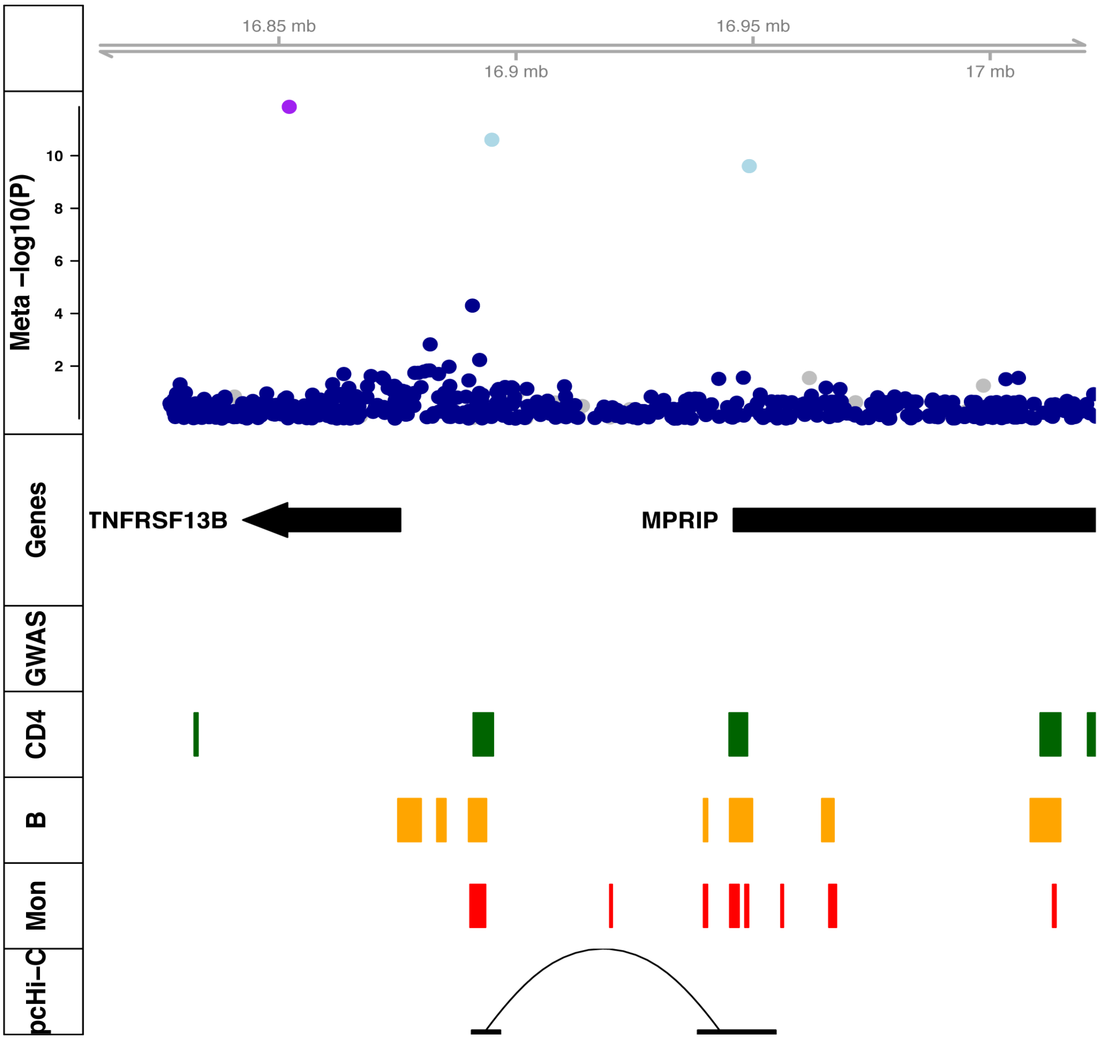


**Figure 6. Regional AD-PID meta-analysis association plot of 17p11.2 (*TNFRSF13B*/TACI*).*** Tracks as described in Figure 3. Tracks are as follows: **Meta -log(P)** dot plot of AD-PID fixed effects association meta-analysis with Li *et al.* (total cases n=1,511; controls n=20,224). The index SNP is purple, others are coloured based on LD information from UK10K project with red indicating high LD (r^2>0.9), blue low LD (r^2<0.2) and grey where no information available. **Gene** - Canonical gene annotation (Ensembl V75), **GWAS** - location of index variants from other immune-mediated disease, **CD4**, **B**, **Mon** - putative regulatory regions in CD+T-cells, Total B cells and Monocytes computed from the union of ATAC-Seq and H3K27ac ChIP-Seq data, **pcHi-C** - Promoter Capture Hi-C interactions, in above primary cell types, with the exception of **Meta -log(P)** which shows dot plot of AD-PID association meta-analysis with Li *et al.*

#### Supplementary Note 4

#### AD-PID GWAS Enrichment Method

We created a compendium of 9 imputed case/control GWAS of European ancestry from sources referenced in the relevant publications (**Table 1**). We next removed the MHC region (chr6:24-45Mb) due to its known association with immune-mediated disease (IMD) and its complex LD structure. We constructed QQ plots for AD-PID GWAS P-values conditioned on strata of association (P < [1,0.1,0.02,$10^{-3}$,$10^{-4}$,$10^{-5}$]) for corresponding variants for each of the nine traits.

| Label | Trait | Cases | Controls | N |
|---|---|---|---|---|
| SLE | Systemic Lupus Erythematosus[1] | 4,036 | 6,959 | 10,995 |
| T1D | Type 1 diabetes[2] | 5,913 | 8,829 | 14,742 |
| CD | Crohn's disease[3] | 12,194 | 28,072 | 40,266 |
| UC | Ulcerative colitis | 12,366 | 33,609 | 45,957 |
| RA | Rheumatoid arthritis[4] | 14,361 | 43,923 | 58,284 |
| Asthma | Asthma[5] | 14,085 | 78,768 | 92,853 |
| T2D | Type 2 diabetes[6] | 266,76 | 132,532 | 159,208 |
| Allergy | Allergy[7] | 96,794 | 145,775 | 242,569 |
| CAD | Coronary artery disease[8] | 345,41 | 261,984 | 296,525 |

**Table 1: Imputed European ancestry GWAS included in enrichment analysis.**

This provided visual evidence for the enrichment of common variants associated with immune-mediated and allergic disease and AD-PID, observed as increasing inflation of AD-PID P-values on conditioning (**Figure 1**).

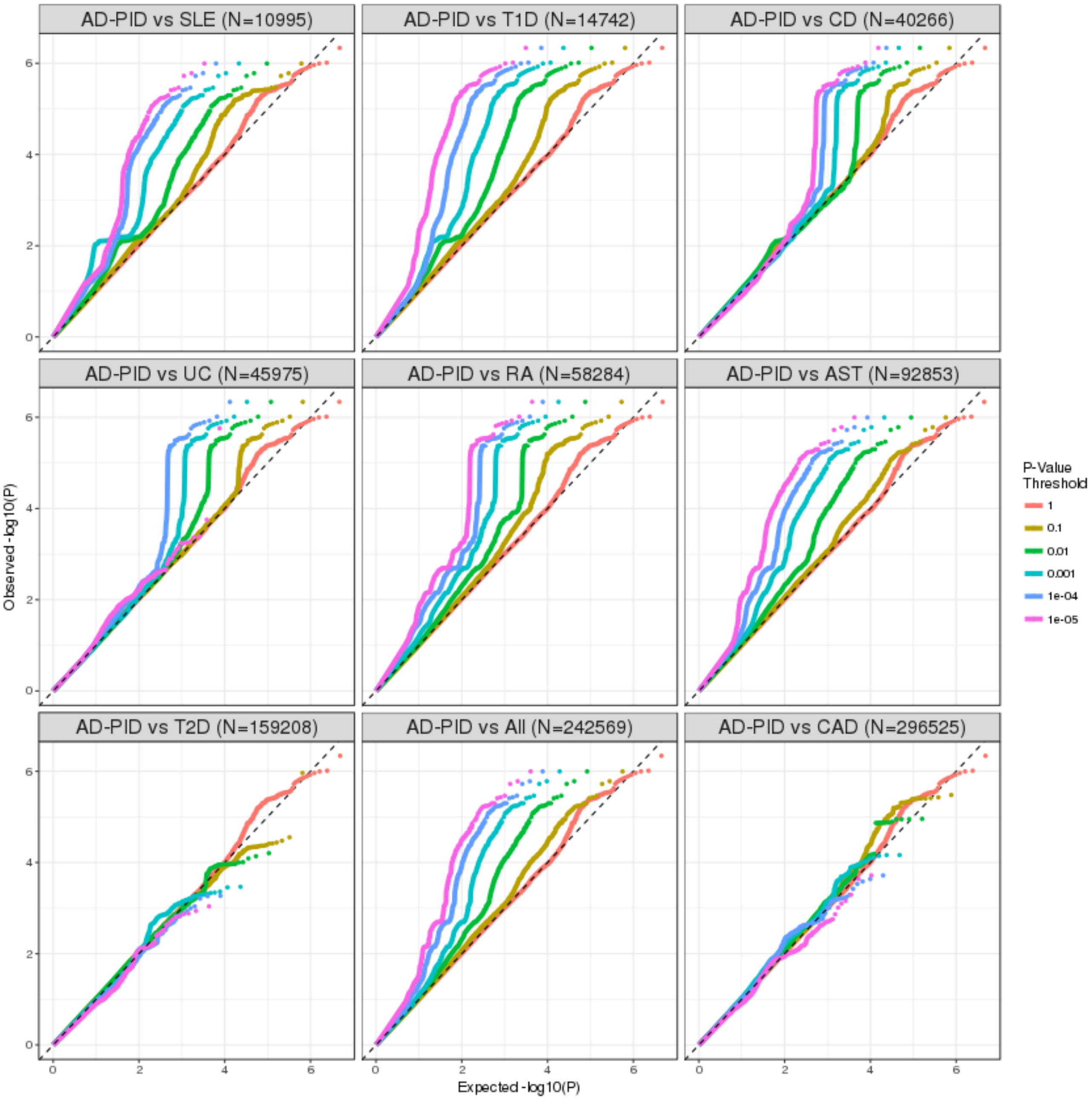


**Figure 1. Conditional QQ plots of AD-PID GWAS with 9 imputed disease case/control GWAS of European ancestry (Table 1)**. SLE – systemic lupus erythematosus, T1D – type 1 diabetes, CD – Crohn's disease, UC – ulcerative colitis, RA – rheumatoid arthritis, AST – asthma, All – Allergy, T2D – type 2 diabetes and CAD – coronary artery disease. N indicates the total sample size of each GWAS AD-PID expected and observed -log10(P) are ploted conditional on subsets of variants stratified by association P-value in the labelled disease. Left deflection of from x=y of curves indicates enrichment of AD-PID GWAS summary statistics in the labelled trait.

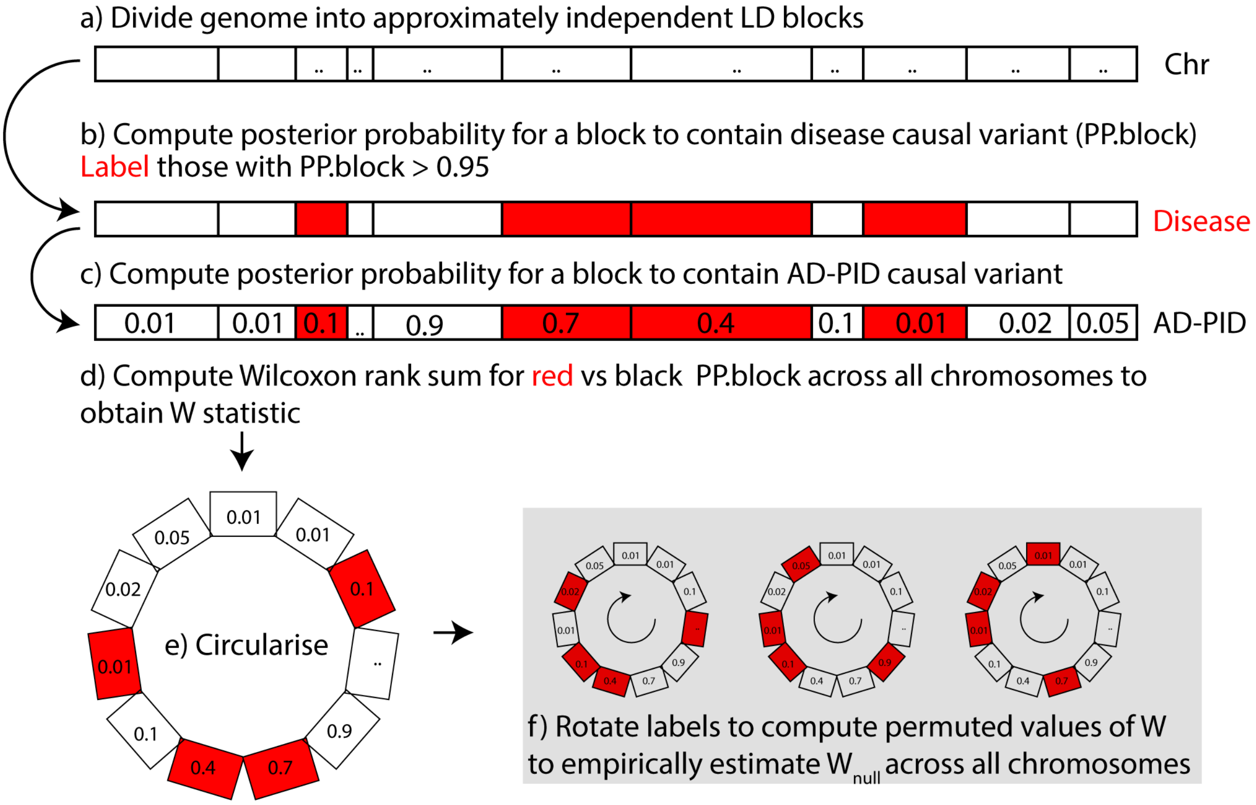


**Figure 2. Schematic of the method to assess enrichment of AD-PID associated variants within the GWAS compendium.** The grey box denotes the permutation step and is carried out multiple times to rapidly compute many estimates for the value of the null Wilcoxon statistic. For a more detailed explanation see text.

Due to the size of the AD-PID cohort, we were unable to use LD-score regression to assess genetic correlation[9]. We therefore adapted a previous method 'blockshifter' [10] in order to assess the enrichment of AD-PID variants with each of the nine traits. For each trait (including AD-PID) we split SNPs into approximately independent LD blocks using HapMap recombination frequencies (**Figure 2a**). We used methods described in [11] and [12] to compute, within a given block, the posterior probability that a given variant is causal (under single causal variant assumptions):

$$\mathrm{sCVPP}_i = \frac{\mathrm{aBF}_i \pi_i}{\sum_{j=1}^{n} (\mathrm{aBF}_j \pi_j) + 1},$$

where $\mathrm{sCVPP}_i$ is the posterior probability that the $i^{th}$ variant in a block is causal, $\mathrm{aBF}_i$ is the asymptotic Bayes factor for the $i^{th}$ variant and $\pi_i$ is the prior probability for the $i^{th}$ variant to be causal. Similarly, j indexes the n variants within the approximately independent LD block under consideration. Here $\pi_i = \pi_j$ are flat prior probabilities for a randomly selected variant to be causal and we use the value $10^{-4}$ used in previous studies employing this method [13]. The benefit of this approach is that it naturally adjusts for LD within a block as sCVPP are modelled jointly across a block.

For each compendium trait we summed sCVPP with each block in order to obtain an overall posterior probability for a block to contain a causal variant (PPCB). We used this to label blocks as to whether there was a high posterior probability (> 0.95) for them to contain a causal variant (**Figure 2b**). We transferred these block labels from the trait under consideration to AD-PID as the block genomic intervals are the same. This operation results in labelled blocks of AD-PID PPCB (**Figure 2c**).

We next used a Wilcoxon rank sum test to evaluate evidence for the enrichment for larger AD-PID PPCB in labelled compared to unlabelled blocks (**Figure 2d**). Such an approach may be confounded by residual LD between blocks and block size. In order to adjust for this we used a circularised permutation technique to compute a suitable null statistic. For a given chromosome, we select all blocks and circularise such that the beginning block adjoins the end of the last (**Figure 2e**). Permutation proceeds by rotating block labels whilst maintaining PPCB across all chromosomes. Whilst these permutations can be computed rapidly, unlike a conventional permutation strategy they also conserve local block labelling structure (**Figure 2f**). We sampled $10^4$ permutations from this pool in order to empirically estimate the mean and variance of the Wilcoxon rank sum statistics under the null (i.e. that there is no enrichment). This was used to generate a standard normal Z score as follows:

$$Z = \frac{W - \bar{W}_{\mathrm{Null}}}{\sqrt{V^*}},$$

where W is the observed Wilcoxon statistic and $W_{\mathrm{Null}}$ and $V^*$ are the mean and variance of the permutation-derived null Wilcoxon statistics. The Wilcoxon and resultant Z scores were computed using the *wgsea* R package [https://github.com/chr1swallace/wgsea].

Whilst GWAS study sample sizes could bias the results of the enrichment analysis, with larger studies likely to find more associations, we did not find evidence for this (**Figure 1** and **Figure 3**). When we plotted enrichment Z scores against underlying GWAS sample size, we observed that CAD and T2D studies that were amongst the largest still showed the least enrichment (**Figure 3**). We are therefore confident that GWAS sample size is not biasing our overarching result of the enrichment of AD-PID associated variants in allergic and immune-mediated disease.

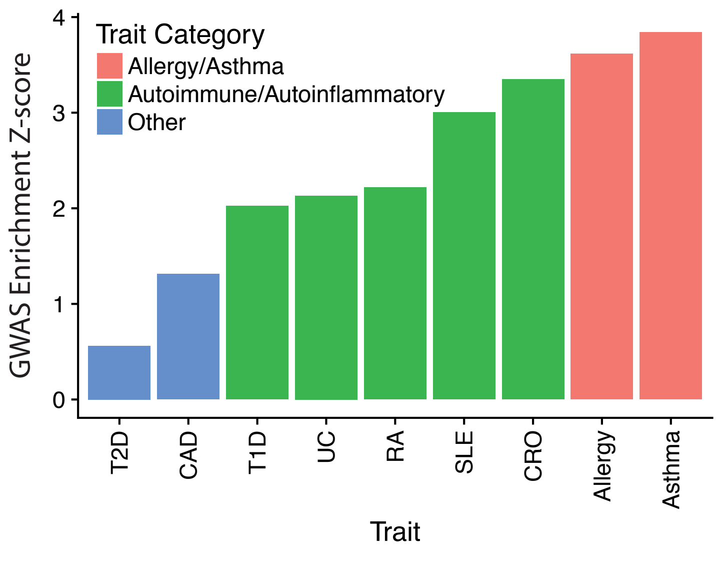


**Figure 3. Immune mediated trait enrichment of AD-PID association signals.** CAD – coronary artery disease, CRO – Crohn's disease, RA – rheumatoid arthritis, SLE – systemic lupus erythematosus, T1D – type 1 diabetes, T2D – type 2 diabetes and UC – ulcerative colitis. The method for generating the GWAS enrichment Z-scores is described in the text.

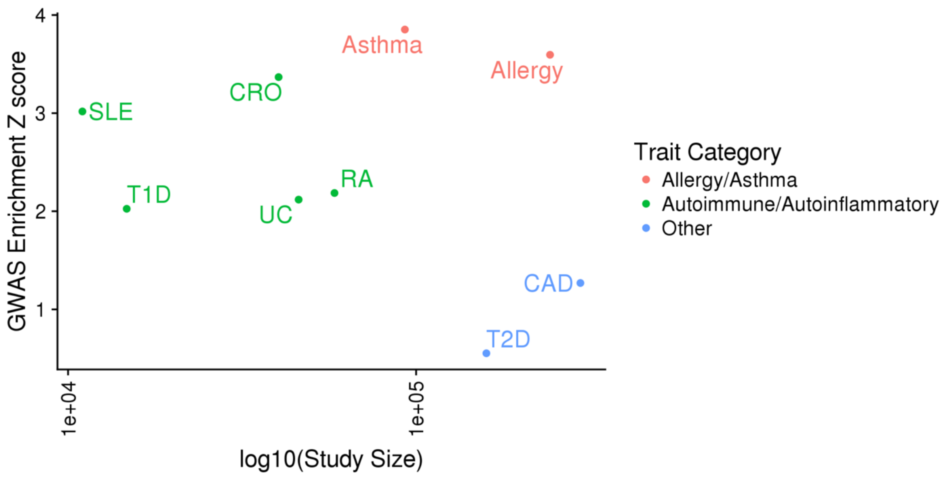


**Figure 4. Scatter plot of AD-PID GWAS enrichment vs overall sample size.** SLE - systemic lupus erythematosus, T1D - type 1 diabetes, CD - Crohn's disease, UC - ulcerative colitis, RA - rheumatoid arthritis, T2D - type 2 diabetes and CAD - coronary artery disease.

### Supplementary Note 5

### PID diagnosis: a comparison of WGS vs WES+SNParray

This paper has demonstrated that a cohort-based whole genome sequencing (WGS) approach can allow increased diagnosis and new gene discovery even in sporadic PID, assessing both rare and common variants. Aspects of this could, in theory, be achieved by a combination of whole exome sequencing (WES) with array-based genotyping and imputation. Here we compare these two possible approaches to PID, focussing on scientific and the cost issues.

#### Scientific comparison

**Common variant genotyping:** Direct sequencing of common variants provides more accurate genotyping than imputation, especially for samples from ethnic groups not well represented by the reference haplotypes (such as those of the 1000Genomes Project) commonly used for imputation.

**Structural variants:** WGS detected single exon and partial gene deletions that are difficult to call by WES CNV algorithms. CNVs generally require higher average coverage to detect by WES, typically 70-80x, compared to the 30-40x to detect by WGS, owing to the read mapping information available in WGS that is used to identify breakpoints. WES, on the other hand, relies only on the relative changes in coverage, and requires across- and within-sample normalisation in order to reliably call changes in copy number. In addition to CNVs, other structural variants such as chromosomal inversions and translocations are more likely to be detected by WGS as the breakpoints usually occur outside the coding regions.

**Detection of pathogenic variants:** Based on the coverage of previously reported PID pathogenic variant sites in aggregated ExAC exomes and our genomes, over 5% of SNVs and InDels have insufficient coverage (<20X) in WES and may therefore be missed by diagnostic WES screening of known genes (**Figure 1**). The figure below shows coverage across all HGMD mutation sites in the known PID genes.

**Additional analyses:** WGS is superior to WES or array for analyses such as autozygosity mapping and detection of uniparental isodisomy (UPD), as it captures all common variants and therefore provides more accurate breakpoints of the affected region. This means that both UPD and unmasked pathogenic homozygous variants can be detected through a single method (WGS) rather than, traditionally, by a genotyping array followed by candidate gene or exome sequencing.

**Non-coding space:** WGS is vital for discovery of defects in the non-coding space, which we have demonstrated can impact on regulatory regions and contribute to pathogenesis. Efforts to identify such defects are becoming more sophisticated, and are increasingly being incorporated into statistical algorithms such as BeviMed. These approaches only work with sufficient statistical power driven by the cohort sample size, and thus as the WGS cohort size grows, our ability to detect defects in the non-coding space will grow with it. A WES approach would not allow exploration of the non-coding space.

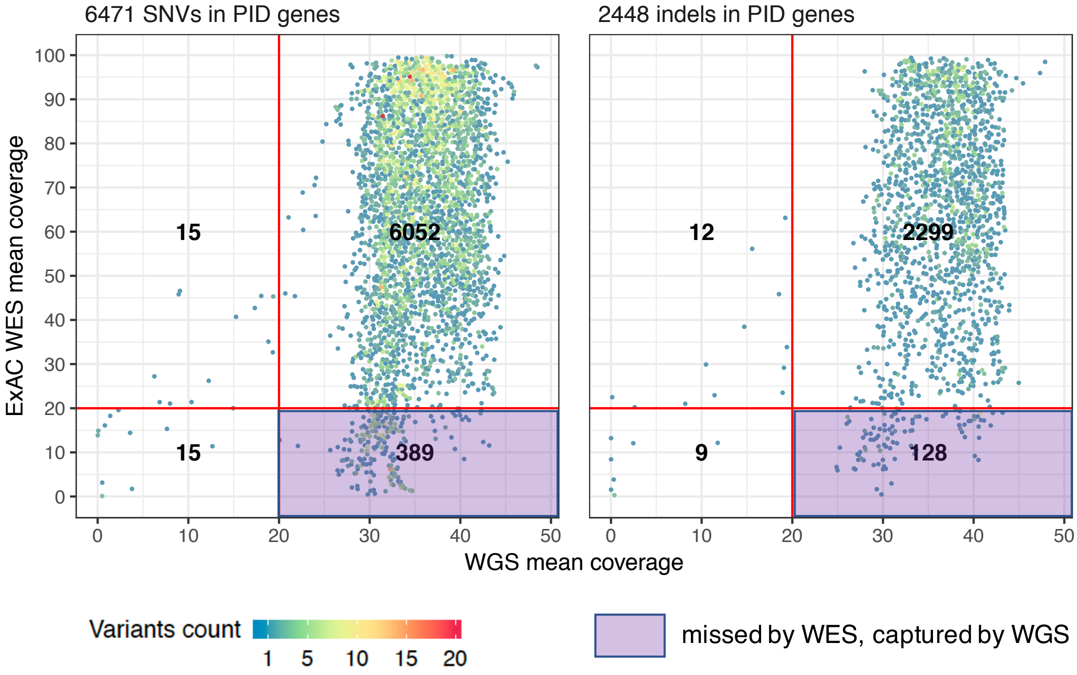


**Figure 1. Comparison of WGS and WES for clinical genetic testing.** WGS and WES coverage of the DM and DM? (denoting disease-causing and probable pathogenic variants, respectively) classes of HGMD SNVs and indels in the IUIS 2017 PID genes. The red axes show the read depth threshold for clinical reporting. The number of variants in each quadrant is indicated, purple – reported by WES only, orange – reported by WGS only, grey – not reported by WES or WGS. For small deletions the coverage was taken as the mean coverage of the deleted bases ± 1bp; for small insertions the coverage was taken as the mean coverage of the two bases surrounding the insertion breakpoint. In line with the metrics downloaded for ExAC, coverage in WGS was obtained using samtools depth (base quality ≥ 10 and mapping quality ≥ 20). The WGS mean coverage was computed on a subset of 1,000 male samples. The WES mean coverage was obtained from ExAC release 0.3.1.

#### Specific findings in this study dependent on WGS

1. The **promoter region of *ARPC1B*** is not targeted by the exome panels used for the ExAC dataset, and 9kb is too small resolution to be reliably called by copy number analysis of SNP arrays, so the pathogenic promoter deletion would have been missed without WGS (see **manuscript Fig. 3c**).

2. WES sensitivity for **single exon deletions** with breakpoints outside the targeted regions is generally poor, as the CNV calling algorithms rely only on relative read depth within and across the samples. WGS, on the other hand, usually has coverage across these non-exonic breakpoints, and the algorithms use read mapping information in addition to relative coverage to make a CNV call, which is a much more sensitive methodology. In our dataset, we detected and reported as pathogenic two single exon deletions (in *CTLA4* and *LRBA*) that would have been unlikely to be called by either WES or genotyping arrays.

3. WGS provided direct **genotyping of rare non-coding variants**, not only allowing for rare variant GWAS that identified genome-wide associations at the *TNFRSF13B* (*TACI*) locus, with two non-coding rare signals in addition to C104R, but future-proofing this dataset for further association discoveries once sample sizes and power increase.

4. **Phasing of compound heterozygous variants** in *DOCK8* was possible because we had WGS coverage between the two variants across the whole gene (coding and intronic regions), which provided a scaffold to build haplotypes using the Nanopore long-range reads (shown in **Extended Data Fig. 5c**). Nanopore sequencing is still highly error-prone and would have required many expensive flowcell runs to generate sufficient data for haplotype calling without relying on the high-quality short-read scaffold.

#### Cost

From a purely cost perspective, once the cost and time (~2 days) of additional lab work required for WES pull-down and separate SNP array experiments are considered, WES+array are currently only marginally cheaper than WGS, and the cost of WGS is still dropping. As an example, before this project commenced in 2012 the list price of a single genome was £5,000. This fell in a step-wise fashion to £650 in 2014. For the next stage of the Genomics England WGS sequencing of 500,000 genomes, the price has dropped to £450 and should reach £200 by 2022. Imputation would also take some additional computing time, and CPU and staff costs. Furthermore, in our experience WGS is easier to manage from an organizational viewpoint than WES+array+imputation, as all the data are generated through a single experiment rather than three.

On balance, we believe WGS already represents a cost-effective approach to PID diagnosis which, provided large enough cohorts are available for analysis, should facititate on-going gene discovery and thus improved diagnostic capacity. Building WGS cohorts could be considered an investment in the future of the diagnosis of PID and other conditions.
